# DeepMOCCA: A pan-cancer prognostic model identifies personalized prognostic markers through graph attention and multi-omics data integration

**DOI:** 10.1101/2021.03.02.433454

**Authors:** Sara Althubaiti, Maxat Kulmanov, Yang Liu, Georgios V Gkoutos, Paul Schofield, Robert Hoehndorf

## Abstract

Combining multiple types of genomic, transcriptional, proteomic, and epigenetic datasets has the potential to reveal biological mechanisms across multiple scales, and may lead to more accurate models for clinical decision support. Developing efficient models that can derive clinical outcomes from high-dimensional data remains problematical; challenges include the integration of multiple types of omics data, inclusion of biological background knowledge, and developing machine learning models that are able to deal with this high dimensionality while having only few samples from which to derive a model. We developed DeepMOCCA, a framework for multi-omics cancer analysis. We combine different types of omics data using biological relations between genes, transcripts, and proteins, combine the multi-omics data with background knowledge in the form of protein–protein interaction networks, and use graph convolution neural networks to exploit this combination of multi-omics data and background knowledge. DeepMOCCA predicts survival time for individual patient samples for 33 cancer types and outperforms most existing survival prediction methods. Moreover, DeepMOCCA includes a graph attention mechanism which prioritizes driver genes and prognostic markers in a patient-specific manner; the attention mechanism can be used to identify drivers and prognostic markers within cohorts and individual patients.

**Author summary:** Linking the features of tumors to a prognosis for the patient is a critical part of managing cancer. Many methods have been applied to this problem but we still lack accurate prognostic markers for many cancers. We now have more information than ever before on the state of the cancer genome, the epigenetic changes in tumors, and gene expression at both RNA and protein levels. Here, we address the question of how this data can be used to predict cancer survival and discover which tumor genes make the greatest contribution to the prognosis in individual tumor samples. We have developed a computational model, DeepMOCCA, that uses artificial neural networks underpinned by a large graph constructed from background knowledge concerning the functional interactions between genes and their products. We show that with our method, DeepMOCCA can predict cancer survival time based entirely on features of the tumor at a cellular and molecular level. The method confirms many existing genes that affect survival but for some cancers suggests new genes, either not implicated in survival before or not known to be important in that particular cancer. The ability to predict the important features in individual tumors provided by our method raises the possibility of personalized therapy based on the gene or network dominating the prognosis for that patient.

## Introduction

Genetic or genomic approaches to understanding disease typically use single or at most a handful of variants within a patient population to identify risk and molecular etiology. However, the phenotypic manifestation of a disease is dependent on the genetic background, which makes elucidation of the causative gene or dysregulated process challenging in complex diseases such as cancer. Even for cancers where there are inherited, penetrant, predisposing germline genetic variants, the outcomes, treatment response characteristics, and prognosis based on single gene or gene panel sequencing can be extremely variable on a patient-to-patient basis [1, 2]. For those cancers where there is no known predisposing variant (arguably the majority), genomic approaches to the discovery of prognostic, predictive or diagnostic markers are often insufficient in themselves to usefully stratify populations and, importantly, to drive personalized approaches to therapy [3, 4]. Consequently, and despite the discovery of cancer driver genes for many cancers and successful implementation of the knowledge that these bring, there is limited success in their translation into clinical useful biomarkers with few cancer prognostic biomarkers currently being approved by regulatory agencies [5–7].

With the advent of high-throughput technologies that capture the physiological landscape of the metabolism, epigenome, RNA and protein expression, and other datatypes, the amount of information available for the identification of new biomarkers and new insights into pathophysiology is increasing almost exponentially [8]. In many ways, the capture of omics knowledge about a single tumor integrates the state of gene expression across the whole genome with that induced by the environment, and increasingly offers a rich and deep picture of the particular state of the cancer cell on a patient-to-patient and population-to-population basis. The addition of fundamental background knowledge, such as cell-type-of-origin [9] and clinical information, to the description of tumor or patient can further enrich the data available for prediction of drug resistance and patient survival. Yet, the challenges of integrating such knowledge, which is often categorical, with quantitative omics data have meant that there are few examples of successful implementation. The combination of multiple types of omics together with other types of data might therefore be expected to facilitate methods that can predict patient-specific outcomes and guide clinical decision-making [10–12].

Several large projects, such as the Cancer Genome Atlas (TCGA) [13], Molecular Taxonomy of Breast Cancer International Consortium (METABRIC) [14], and TARGET [15], have characterized different types of cancer on multiple levels and generated different types of omics datasets for these cancers. We exploit this rich data and integrate it with background knowledge to develop a model for cancer survival, and to highlight genes that make the most significant contribution to the model on a patient-by-patient basis.

There are several different methods available for integration and analysis of multi-omics data [16, 17]. One of the key challenges is the high dimensionality of data which can be addressed through unsupervised machine learning to generate latent, lower-dimensional representations that are subsequently used for prediction tasks [18, 19]. These methods may also allow incorporation of background knowledge such as pathways or biological interaction networks which are crucial for understanding and representing cancer pathophysiology [20, 21]. There are now a large number of methods for machine learning with multi-omics data [12, 22–24] using a wide range of different approaches; a common approach is the prediction of survival time for which benchmark datasets have been developed [25].

To model cancer progression, quantitative relationships and dependencies between functional elements within a cell need to be captured. While these dependencies are traditionally characterized using systems biology models based on ordinary or partial differential equations [26], these quantitative relations are not immediately accessible in many machine learning models. Furthermore, while there has been significant progress in interpretability of machine learning methods [27], they are not yet regularly applied to high-dimensional biological data.

We developed DeepMOCCA, a computational model using graph convolutional neural networks that incorporates background knowledge in the form of interaction networks. DeepMOCCA is an end-to-end deep learning model that predicts survival from cancer multi-omics data and generates a representation of nodes and cancer samples; using an attention mechanism, DeepMOCCA can identify cancer drivers and prognostic markers in individual samples and stratify cohorts based on molecular characteristics. To ensure that DeepMOCCA focuses primarily on the dynamic interactions that occur within a cell, it relies only on information about the sample, i.e., omics data and the tumor type and anatomical location, but does not incorporate clinical information (e.g., age or sex) which may correlate with cancer progression and survival but does not provide information on molecular or cellular patho-etiology. We illustrate that DeepMOCCA predicts survival time accurately and similarly to competing methods (including those that incorporate clinical information) and reliably identifies cancer drivers and prognostic markers. We also make DeepMOCCA freely available as a software tool, including all the necessary steps to train the model, so that it can be adapted easily to related applications.

## Results

### Integration and analysis of multi-omics data in graph neural networks

We developed a machine learning approach to learn representations of the multi-scale activities and interactions within a tumor from multi-omics data associated with individual cancer samples by predicting an easily obtainable measure, the survival time. Our model takes as input data derived from individual samples, in particular the set of germline and somatic variants, absolute methylation in normal and tumor tissue, absolute gene expression in normal and tumor tissue, copy number variants detected in tumor tissue, and the cancer type and anatomical location. We use this information to calculate differential expression and differential methylation and determine the cell type of origin.

Our approach leverages background knowledge to address three key challenges: integration of different types of omics data; modeling the dynamics and interactions within a cell; and interpretation and explanation of the analysis results. We integrate the different types of data using biological background knowledge in the form of a graph in which nodes represent genes, transcripts, and proteins, and edges between nodes represent (genetic or physical) interactions between them. For this purpose, we design a set of mapping functions that map the information from the multi-omics data to nodes in this graph. Using genetic variants in germline and somatic genomes, we assign a value to gene nodes that represents the pathogenicity prediction score for the most pathogenic variant within that gene; if a variant is intergenic, we assign its pathogenicity score to the nearest gene. For absolute gene expression in the tumor, we assign the absolute expression value of a transcript to the node representing that transcript. Differential gene expression and differential methylation are each used to assign a single value to each node based on the fold change between normal and tumor tissue for differential expression and the *p*-value of the differential methylation. Copy numbers are assigned qualitatively to gene nodes depending on whether a gene is affected by a deletion or duplication.

As a result, we obtain a graph in which nodes are assigned a list of values for each sample. Some samples lack a particular datatype in which case we treat the values as missing. The edges between nodes in the graph represent functional interactions. We hypothesize that some of the omics features we include (or the combination of features) localize on this graph, i.e., that the attributes of nodes in small connected subgraphs are significantly related to observable phenotypes. Graph convolutional neural networks can exploit this locality using methods such as message passing between adjacent nodes [28], whereas message passing in our labeled graph corresponds to modeling the affect that features (e.g., gene expression, methylation, or variants) associated with one node have on related nodes. Being able to quantitatively represent and compute these dependencies will capture some aspects of dynamic interactions that occur within a cell.

We use the graph labeled with values derived from an individual samples’ omics data to predict patient survival time using a graph convolutional neural network combined with Cox regression. Cox regression is a means to account for censored data in regression analysis; integrating Cox regression with a graph convolutional neural network allows us to train the model in an end-to-end fashion to predict survival time in individuals from the samples’ omics data. As our graph is based on functional interactions between genes or proteins and uses message passing to generate node representations, back-propagation used during training will generate the quantitative dependencies between nodes. Figure 1 shows the model we use.

**Fig 1.**
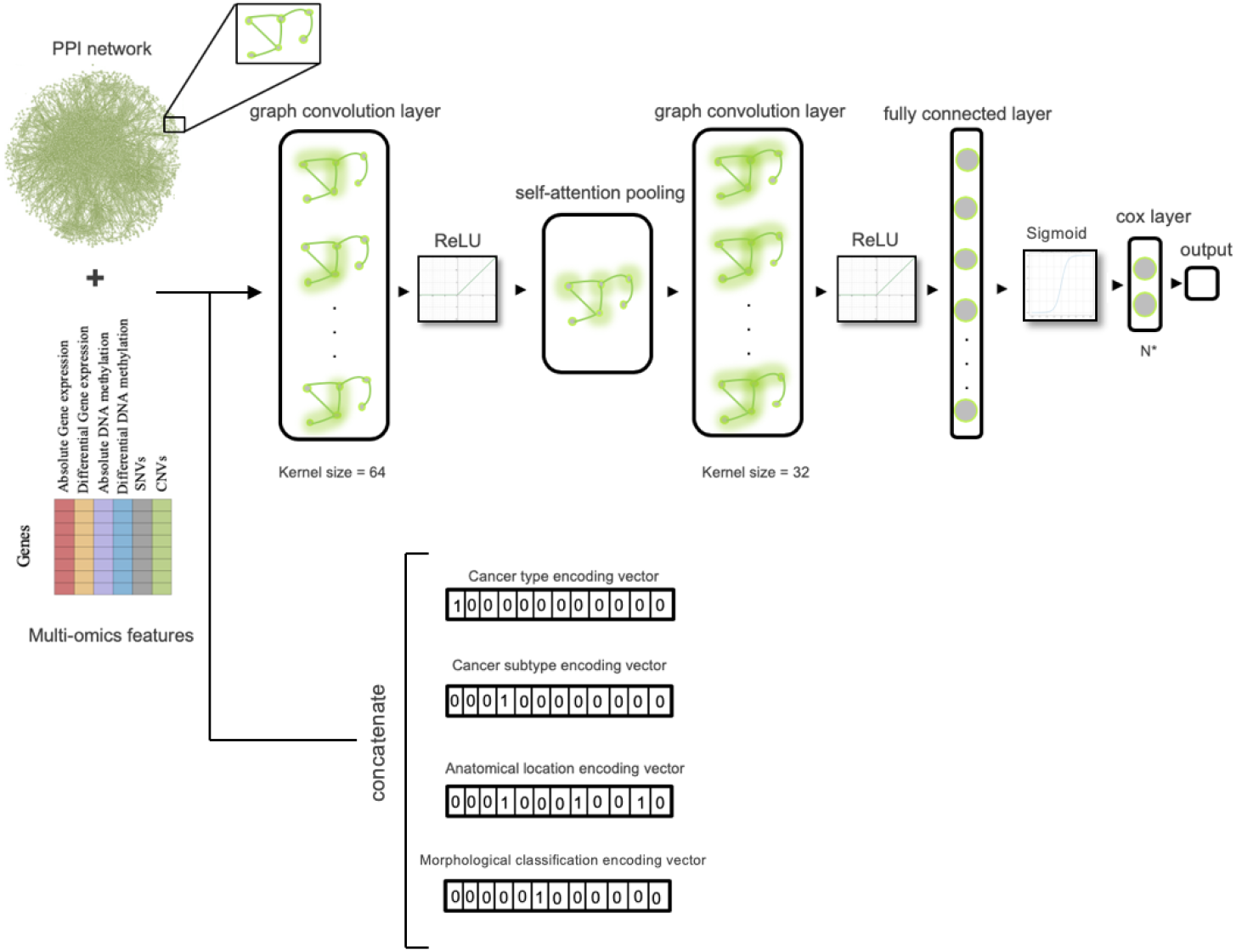
Graph convolutional neural network model for survival time prediction across different cancer types.

Initially, we apply our model to breast cancer, lung cancer, and glioblastoma cohorts so we can evaluate the impact of different types of omics data on survival time prediction on a small set of different cancer types. We evaluate model performance using the Concordance index (C-index) to measure the difference between predicted and assigned survival time. Results are summarized in Supplementary Table 1. We find that, individually, differential methylation, absolute methylation, and differential gene expression are most predictive of survival time. Combining different omics features provides significant improvements in predicting survival time, with the highest performance achieved when all types of omics data are combined.

Our model also allows us to test different ways of representing omics data. We tested different ways to normalize values assigned to genes as these normalizations convey different biological information; in the matrix of values assigned to genes from cancer samples, we can normalize values across the entire matrix, across each row (cancer sample), or across each column (gene). While a global normalization is more common, row-based normalization allows us to highlight values that are significantly higher or lower within one sample (e.g., which genes are expressed at high or low levels within a single sample), and column-based normalization allows us to highlight values assigned to a particular gene that are significantly higher or lower within one sample (e.g., whether a gene is expressed at higher or lower levels within one sample compared to all others). We find that column-based normalization performs better than row-based normalization, while the global normalization approach performs close to random. The best results are achieved when combining both row- and column-based normalization (Supplementary Table 2).

After optimizing our model on these three cancer types we developed a joint model that can predict survival time for 33 types of cancer. While a main motivation of having a joint model for multiple types of cancer is simplicity, we also tested whether information can be transferred between different cancer types and therefore improve overall predictive performance (Supplementary Table 4). To allow our model to distinguish between multiple types of cancer, we add the clinical cancer type as well as the anatomical site of the cancer as additional inputs; furthermore, based on the cancer type we assign the cancer’s morphological classification and use it as another input. All this additional information is routinely obtained through clinical investigations and available for any tumor sample. Performance results for evaluating the model are shown in Supplementary Table 3. We find that the joint model can further improve over models trained on only single cancer types, showing that at least some transfer of information occurs when combining information from different cancer types. We also use this model to compare our performance to other efforts to predict survival from cancer multi-omics data. Other models generally predict survival time only for single types of cancer and use additional features beyond omics data, including images. We individually compare our model to other models predicting survival time on 23 different cancer types, and find that our joint model improves over predictive performance observed in other models on the same datasets (see Table 1). In addition to our deep learning model architecture, the main difference between related methods and DeepMOCCA is the use of an interaction network, indicating that use of protein interactions as background knowledge can improve cancer survival analysis.

**Table 1.**
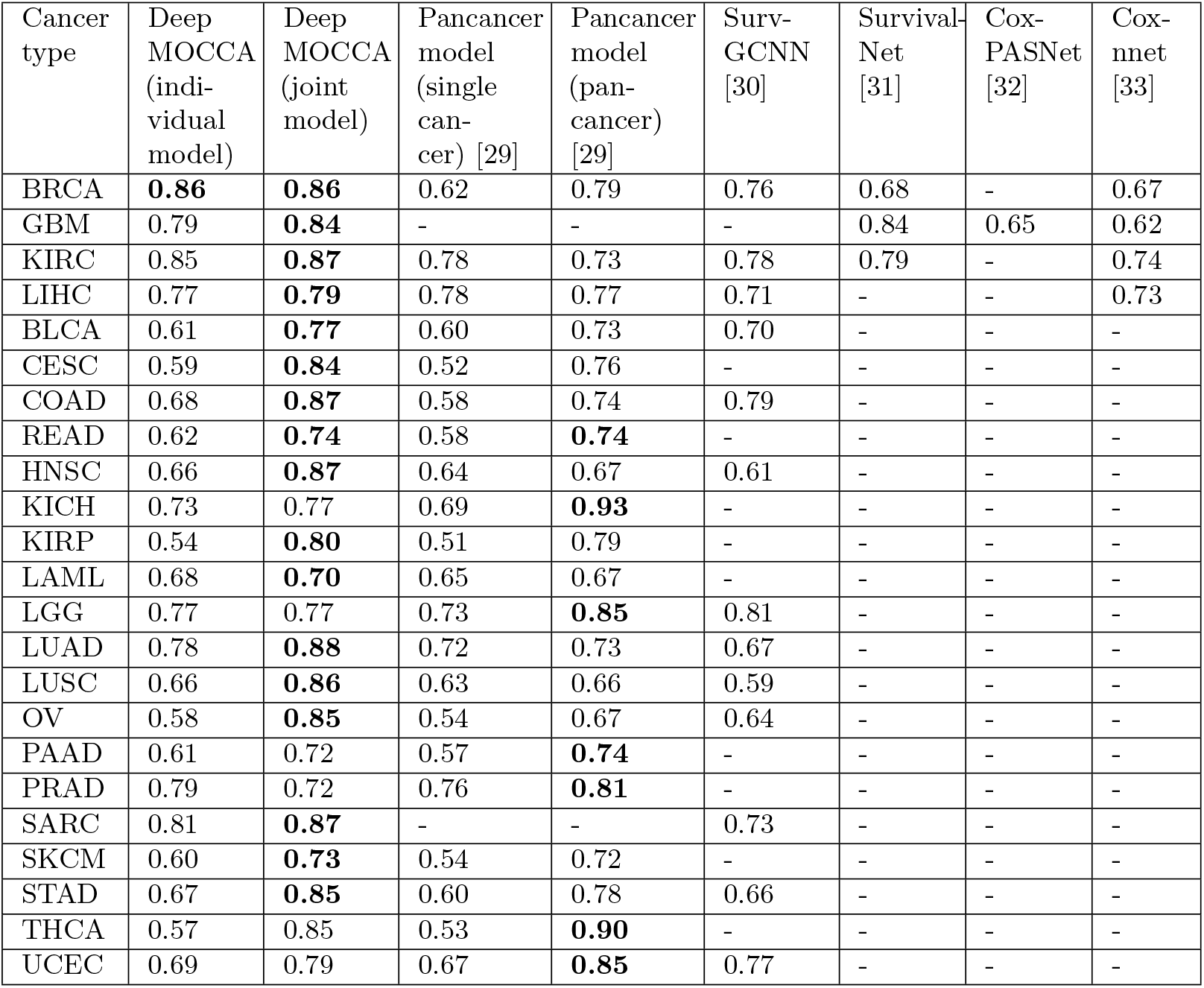
Performance comparison based on the C-index. For the glioblastoma dataset, SurvivalNet combines the GBM and LGG datasets while we treat them separately. For COAD and READ datasets, [29] combines both and we report their performance measure twice in both datasets. Bold numbers indicate the best performance obtained across the set of algorithms.

### Exploring graph attention

We include an attention mechanism in our model which allows us to identify the graph nodes that are important for predicting survival. In contrast to methods such as feature selection or ranking, graph attention provides a mechanisms that identifies feature importance in a sample-specific manner. This mechanism therefore allows us to rank the importance of graph nodes (genes or their products) for each sample.

For each sample in our evaluation set, we rank the graph nodes using the attention mechanism. We hypothesize that the highest-ranked nodes represent prognostic markers and cancer drivers. As shown in Table 2, we find that between 96.9% (in uveal melanoma) and 54.1% (in kidney chromophobe renal cell carcinoma) samples have a cancer driver gene listed in COSMIC at the highest-rank for that specific cancer type. Nodes ranked by the attention mechanism within the top ten also represent cancer drivers (between 72.5% for kidney renal clear cell carcinoma and 28.5% for pheochromocytoma and paraganglioma). Furthermore, we find that the attention mechanism also ranks prognostic markers in the highest ranks; average ranks for known prognostic markers in each cancer type are shown in Supplementary Table 5. These results also allow us to identify candidate genes not known to be cancer drivers or prognostic markers but ranking highly across multiple samples (Supplementary File 1).

**Table 2.** Evaluation results for rank-based attention mechanism based on the joint model with respect to the precision at different ranks in identifying known cancer drivers.

| Cancer type | Number of samples | Actual total number of driver genes | ROCAUC | Precision@1 | Precision@5 | Precision@10 |
| --- | --- | --- | --- | --- | --- | --- |
| TCGA-ACC | 80 | 15 | 0.7836 | 0.7595 | 0.3472 | 0.2947 |
| TCGA-BLCA | 407 | 78 | 0.8462 | 0.8747 | 0.6532 | 0.5327 |
| TCGA-BRCA | 1044 | 99 | 0.8608 | 0.8916 | 0.7261 | 0.6049 |
| TCGA-CESC | 294 | 45 | 0.7643 | 0.7285 | 0.5423 | 0.4726 |
| TCGA-CHOL | 36 | 45 | 0.8726 | 0.6648 | 0.4075 | 0.3673 |
| TCGA-COAD | 433 | 45 | 0.8264 | 0.9053 | 0.8159 | 0.6512 |
| TCGA-DLBC | 37 | 85 | 0.7921 | 0.7356 | 0.5093 | 0.4302 |
| TCGA-ESCA | 184 | 71 | 0.8165 | 0.7790 | 0.4487 | 0.3836 |
| TCGA-GBM | 166 | 35 | 0.8644 | 0.6362 | 0.5170 | 0.3528 |
| TCGA-HNSC | 510 | 62 | 0.7637 | 0.5735 | 0.4774 | 0.3647 |
| TCGA-KICH | 66 | 7 | 0.6574 | 0.5406 | 0.4187 | 0.3911 |
| TCGA-KIRC | 339 | 22 | 0.8820 | 0.9175 | 0.8673 | 0.7247 |
| TCGA-KIRP | 288 | 24 | 0.8017 | 0.7549 | 0.6529 | 0.5563 |
| TCGA-LAML | 140 | 61 | 0.8537 | 0.6842 | 0.4914 | 0.3159 |
| TCGA-LGG | 511 | 38 | 0.7758 | 0.8754 | 0.6850 | 0.4630 |
| TCGA-LIHC | 371 | 31 | 0.7681 | 0.7961 | 0.5368 | 0.3546 |
| TCGA-LUAD | 509 | 42 | 0.9039 | 0.8623 | 0.6846 | 0.5295 |
| TCGA-LUSC | 496 | 60 | 0.8378 | 0.8024 | 0.4774 | 0.3305 |
| TCGA-MESO | 83 | 17 | 0.6346 | 0.6735 | 0.4672 | 0.2964 |
| TCGA-OV | 443 | 37 | 0.7963 | 0.6382 | 0.4290 | 0.3151 |
| TCGA-PAAD | 178 | 52 | 0.7952 | 0.7382 | 0.4955 | 0.3080 |
| TCGA-PCPG | 179 | 9 | 0.6119 | 0.5569 | 0.3148 | 0.2846 |
| TCGA-PRAD | 498 | 82 | 0.8474 | 0.8642 | 0.6093 | 0.4523 |
| TCGA-READ | 158 | 72 | 0.9106 | 0.9085 | 0.7536 | 0.4239 |
| TCGA-SARC | 255 | 8 | 0.6846 | 0.7530 | 0.5492 | 0.3751 |
| TCGA-SKCM | 456 | 39 | 0.7551 | 0.7734 | 0.5854 | 0.3482 |
| TCGA-STAD | 414 | 35 | 0.7323 | 0.7915 | 0.6242 | 0.4678 |
| TCGA-TGCT | 134 | 9 | 0.6670 | 0.7891 | 0.5726 | 0.3586 |
| TCGA-THCA | 496 | 40 | 0.8367 | 0.8143 | 0.6480 | 0.5732 |
| TCGA-THYM | 123 | 11 | 0.6994 | 0.9472 | 0.8128 | 0.4307 |
| TCGA-UCEC | 542 | 66 | 0.7580 | 0.8538 | 0.6349 | 0.4287 |
| TCGA-UCS | 55 | 21 | 0.7215 | 0.7154 | 0.5640 | 0.3889 |
| TCGA-UVM | 80 | 11 | 0.6862 | 0.9688 | 0.8525 | 0.5763 |

Genes that are ranked significantly higher by our model’s graph attention mechanism across all samples within a cohort are in Supplementary File 2 (t-test, *α* = 0.05, Benjamini-Hochberg correction). We identify the known driver genes as being ranked significantly higher within their cohorts.

The inputs of our model are omics data derived from individual samples; before our model performs regression for survival time, it generates a “representation” of the input features. These representations may be useful for patient cohort stratification. We illustrate the distribution of the features representation for each cancer patient with a t-SNE visualization in Figure 2.

**Fig 2.**
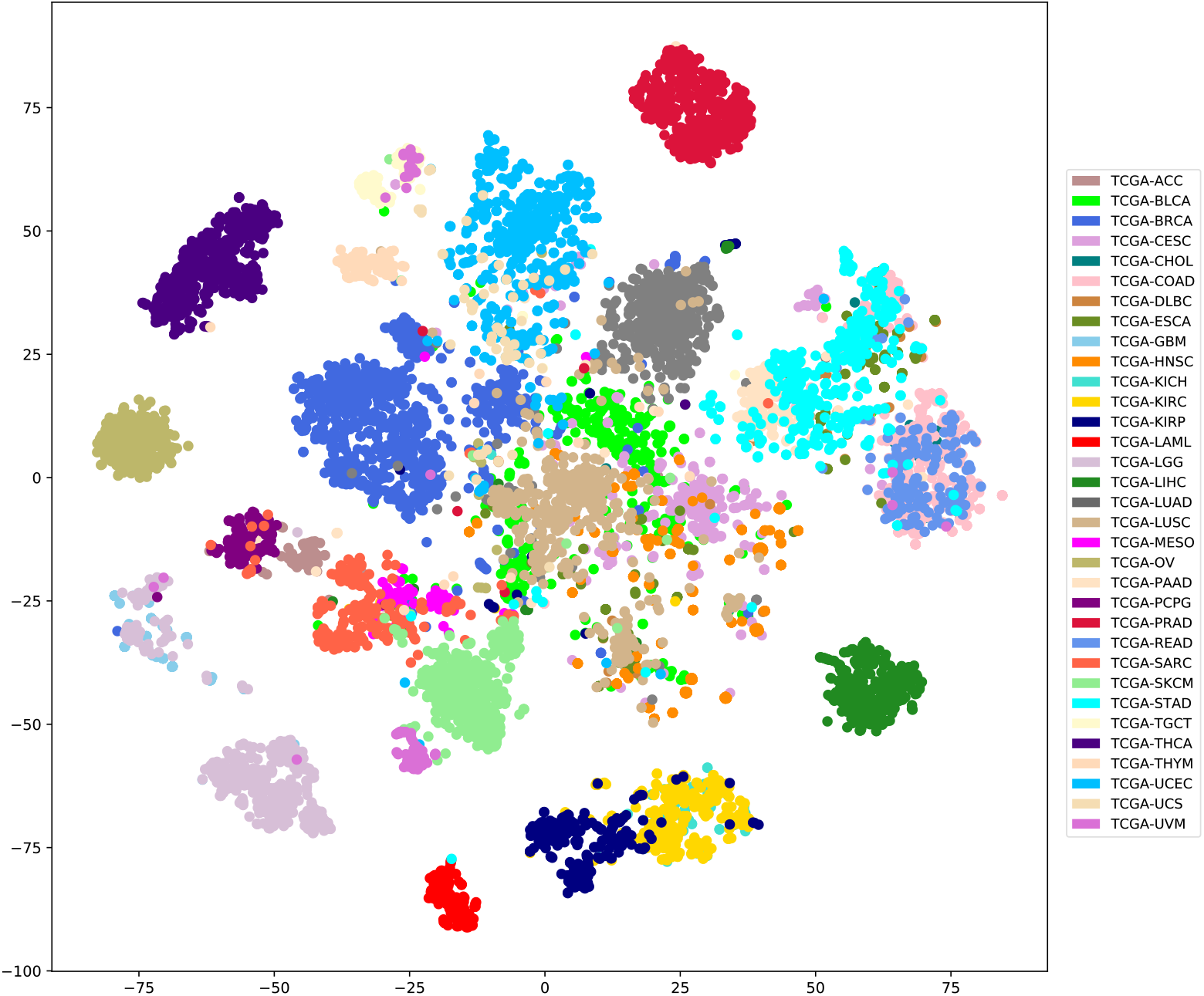
Visualization of features representation using the t-SNE method over the 33 cancer types.

To estimate global similarity between the 33 cancer types included in our study, we identify the pairwise similarity between the average of the generated representations for the 33 cancer types (Figure 3). We find that some cancer types are very similar to other types, such as colon adenocarcinoma (COAD) and rectum adenocarcinoma (READ), or low grade glioma (LGG) and glioblastoma (GBM). The renal tumors kidney renal clear cell carcinoma and kidney renal papillary cell carcinoma (KIRC and KIRP) are similar to each other, but both are also related to prostate adenocarcinoma (PRAD). While this is somewhat unexpected, it is noteworthy that very rare cases of primary renal-type clear cell carcinoma have been described in the prostate [34] and the ontogenic relations of prostate and metanephros are very close. The similarities between ovarian carcinoma and thyroid carcinoma may be unintuitive; however, both cancer types fall into the same subclass (C7) of cancers characterized using multiple omics parameters in a previous study [35] and are cancers with an intermediate prognosis. The representations we use for clustering are based on the prediction of survival and not necessarily linked to tissue-specific oncogenic processes; consequently, we may identify relations between tumors that extend beyond tissue of origin or histological type.

**Fig 3.**
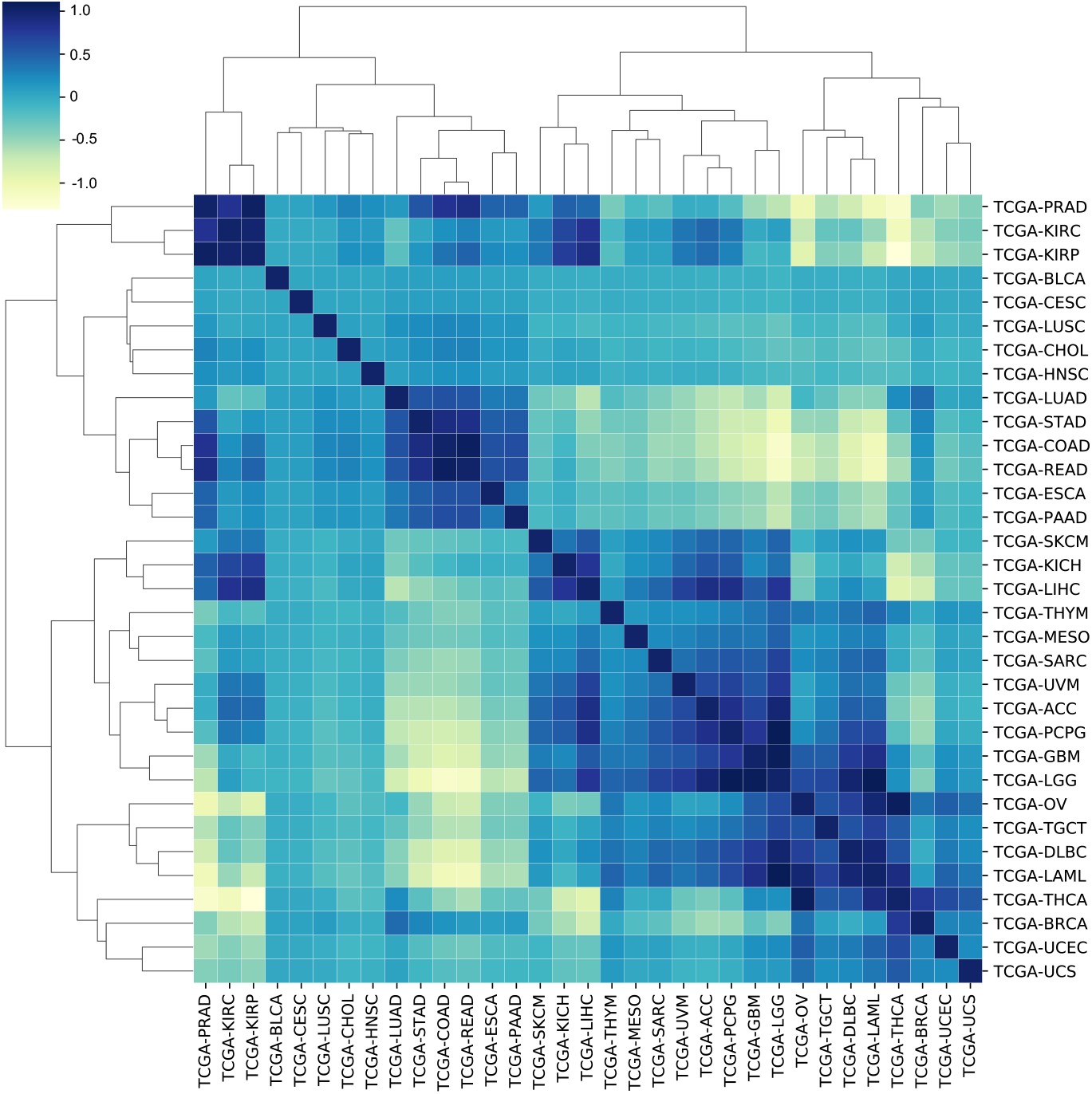
Visualization of the correlation matrix for the 33 cancer types.

## Discussion

### Deep learning on multi-omics data

DeepMOCCA is a method for integrating and analyzing omics data with respect to background knowledge in the form of a graph. While we applied DeepMOCCA to cancer survival prediction, our method can be applied to other phenotypes in which single or multiple types of omics data are available, phenotypes are likely associated with modules of interacting genes or proteins, and for which training data such as survival time is available. Predicting survival time, while useful, is not the main outcome of our work; instead, following a deep learning approach [36], our model generates representations of samples derived from a combination of molecular features and molecular interactions; these representations can be used – in a patient-specific manner – to reveal pathophysiology, cancer drivers, and prognostic markers.

While DeepMOCCA uses interaction networks, other types of background knowledge can be used in a similar manner. The main conditions are that the background knowledge used can be represented as a graph, that the relations convey biologically meaningful information, and that the phenotype correlates with modules (i.e., sets of related nodes) in the graph. Representing knowledge in a graph-based form is increasingly common and forms the foundation of a growing number of bioinformatics resources [37], thereby allowing our method to be used for other types of data and conditions; in the future, we expect similar methods to be adopted using either different types of features or different labels, including quantitative and binary traits.

DeepMOCCA is able to outperform other survival models on most cancer types (Table 1). Similarly to DeepMOCCA, other models have previously explored survival models based on Cox regression and trained using TCGA data [31, 38–41], integrated and utilized multi-omics data for predicting survival or driver genes [32, 42–44], applied transfer learning to develop pancancer models [29], and utilized graph convolutional neural networks to include background knowledge in these models [45]. The majority of prognostic machine learning models include different types of clinical information as part of predicting survival; in particular, using age and sex as covariates can significantly improve predictive performance. DeepMOCCA is different from prior work in several aspects; first, instead of focusing on prognosis as a main goal, DeepMOCCA includes a graph attention mechanism to identify graph nodes (corresponding to genes or proteins) that contribute to the prognosis; DeepMOCCA uses only information derived directly from the tumor (multi-omics data, tumor type, and anatomical location) and specifically highlights the pathobiological mechanisms underlying a prognostic prediction; DeepMOCCA also includes as background knowledge functional interactions between proteins [46] but does not include graphs in which edges have no functional biological interpretation (such as similarity networks). Furthermore, as DeepMOCCA relies primarily on multi-omics data, we spent significant effort to explore different ways to represent and normalize these data types so that the data can meaningfully be used as part of the model optimization process; this likely explains why our model can provide predictions as accurate as other deep learning models that use the same dataset and (transfer) learning approach as DeepMOCCA but also include clinical data (whereas DeepMOCCA does not).

One key limitation of the DeepMOCCA model is the lack of model interpretability with respect to the multi-omics features relevant for a prediction. The attention mechanism in DeepMOCCA outputs graph nodes that are relevant for predicting survival but not the node feature that contributed to the prediction. In the future, an additional mechanism could be added to also highlight the feature of a graph node that contributes to a prediction.

### Identifying prognostic markers and cancer drivers through graph attention

DeepMOCCA can identify prognostic markers and cancer drivers through graph attention (Table 2). In our analysis, most of the markers and drivers we identify in the highest ranks are already well-known.

We used two strategies for further identifying candidate genes from our attention mechanism. The first is at the level of the complete cohort by determining whether a gene is ranked significantly higher than expected through the attention mechanism, and determining the effect size. The second is to examine the frequency with which the gene is implicated at an individual level, i.e., in how many tumors the gene is highly-ranked by attention. The former assumes that the cohort of patients is coherent and unstratified, whereas the latter assigns individual patients to categories. Larger patient groups would improve the detection of subgroups within the overall cohort but we believe that the size of the cohort makes the cohort-wide approach unlikely to support the discovery of novel candidate genes that characterize small cancer subgroups. The identification of candidate genes through graph attention, however, is applicable to single patient samples, and can identify the nodes that are most active in computing survival time for this sample. In both the individual and cohort approaches, DeepMOCCA relies not only on information from mutations alone but also on gene expression and methylation.

We focus here on two cancers with complementary patterns of identified prognostic genes in more detail, breast adenocarcinoma (BRCA) and prostate adenocarcinoma (PRAD). In PRAD, we find that most genes we identify through graph attention have already been identified as driver genes (74 out of 82 genes in the IntOGen database) but we fail to find some well-established prognostic genes (such as *AR, TP53*, and *SPOP*) as being ranked significantly higher across the whole cohort. In some cases, there are possible explanations for this. For example, lack of significance for *AR* in our list likely reflects that the tumors were all primary pre-treatment cases and androgen receptor amplifications and mutations are much more common in treated metastatic tumors than primary tumors [47]. However, these genes are ranked top in several samples (six times for *AR*, seven times for *TP53*, and four times for *SPOP*). For example, somatic mutations in *SPOP* account for 13.7% of tested tumors as estimated by TCGA (data from Genomic Data Commons Data Portal [48]) and 11.0% as estimated by IntOGen [49]. Comparing these frequencies to those we find indicated by graph attention suggests that *SPOP* mutations are not primarily driving survival prognosis in our model. Other previously established markers may also be missed because attention ranks genes specifically based on on their impact on survival time, which in many cases may show a different dependency than tumor initiation, and relates in a complex way to tumor propagation [50]. This hypothesis is supported by direct epidemiological evidence where *SPOP* mutations have been reported not to be associated with cancer-specific survival in the absence of other clinical data [47] (which DeepMOCCA also does not use). Interestingly, also using the TCGA dataset, not even the relationship between favorable histology and *SPOP* mutations in tumors reached significance in another study [51]. It is also interesting that we find *TP53* to be significantly associated with only 13 out of 33 cancer types when testing significance within a cohort; similarly, for most cancer types examined in Donehower et al. [52], no statistically significant differences in overall survival were observed for mutant versus wild-type *TP53* cancers, leading to the conclusion that *TP53* is only useful as a prognostic marker in some contexts.

Our frequency analysis suggests several genes not included in COSMIC or IntOGen that might be candidates for further study. *ZP4*, a component of the oocyte zona pellucida, has not been implicated as a driver gene or in the prognosis of any cancer but ranks first in four samples. *ZP4* has recently been detected in a prostate cell line and prostatic adenocarcinoma with the suggestion that it might be useful as a prognostic marker or be part of tumor immunosurveillance processes [53]. *CRX* ranks first in four samples; *CRX* is a cone-rod homoeobox transcription factor which plays a role in the differentiation program of photoreceptor cells and has been strongly implicated in the growth and differentiation of an aggressive sub-class of medulloblastoma [54]. While still not formally assigned as a cancer driver or prognostic marker, prior work and our findings suggest that *CRX* may predict outcomes in prostate cancer as well as medulloblastoma. The gene is normally expressed at low levels in the prostate, mainly in epithelial cells, where its function is unknown [55]. We also identify *TUBB3* as highest-ranked gene in three samples. *TUBB3* is a beta tubulin gene. So far, it has not been associated as a cancer driver but there is evidence that it is not only a prognostic marker for prostate adenocarcinoma but also indicates responsiveness to docetaxel and is a candidate theranostic marker [56, 57]. Our evidence from graph attention supports this line of investigation and illustrates the utility of our prognostic model.

The BRCA cohort has 1,044 samples and we identify all of the BRCA-associated genes in IntOGen as ranking first in some samples (the 99 driver genes for BRCA listed in IntOGen rank first in between three and 21 samples). However, the statistical power of the cohort-wide analysis only allows us to identify 7 genes as significant. We identify several candidate genes ranked first by attention in multiple patients, such as *TMEM88* and *SDR42E2. TMEM88* is a small transmembrane protein that inhibits Wnt signaling and has been implicated in several cancers, of which breast adenocarcinoma is one [58, 59], but it has not been formally identified as a driver gene (although COSMIC records an effect on chemotherapy drug sensitivity). However, cytosolic *TMEM88* has been correlated with advanced stage and metastasis, and has been proposed as a biomarker in BRCA, ovarian cancer, and non-small-cell lung cancer [59]. Because of the effect of *TMEM88* on the Wnt signalling pathway, identification of *TMEM88* by graph attention in these two samples might indicate that Wnt pathway-modifying treatments might be an effective personalized therapeutic strategy; the low frequency of *TMEM88*-associated samples shows how DeepMOCCA may improve the precision of deciding on personalized treatments.

*SDR42E1* is a short chain dehydrogenase/reductase family member and metabolizes steroid hormones [60]. It is not listed as a driver or prognostic gene in COSMIC or IntOGen. Very little is known about the function of this gene in cancer, but its epigenetic control in colorectal cancer has come under scrutiny [61]. In colorectal cancer, it shows aberrant methylation marks as well as an unusual response to 5-aza-dC in cell lines [62]. DeepMOCCA uses DNA methylation as a feature in modeling survival and it is possible that the epigenetic behavior of *SDR42E1* explains its graph attention score in these samples. While these may be possible explanations, the absence of interpretability on the level of omics features (in contrast to genes) emphasises the need to further develop the DeepMOCCA model and add additional mechanisms that can also identify the specific node features contributing to a prediction.

## Conclusions

DeepMOCCA is a computational model based on machine learning that addresses three challenges in understanding molecular cancer pathobiology: DeepMOCCA integrates multiple type of omics data and background knowledge using a graph-based approach; it predicts survival time in a patient-specific manner using a graph neural network; and it can be interpreted through the use of graph attention. In particular the interpretability of the model, and its application to individual samples (in contrast to cohorts) allows it to be applied as a tool for precision medicine. DeepMOCCA is available as Free Software [63] at https://github.com/bio-ontology-research-group/DeepMOCCA.

## Methods

### Multi-omics dataset

We utilized multiple types of omics data downloaded on 18 May 2020 from The Cancer Genome Atlas (TCGA, http://cancergenome.nih.gov;dbGaPphs000178) [13]. For each type of cancer, we use the data related to gene expression, DNA methylation, copy number variation (CNV), single nucleotide variation (SNV), and associated clinical data. In total, we obtain and use information for 10,005 samples from 33 cancer types; gene expression data is available for 10,558 samples, DNA methylation for 10,943 samples, CNVs for 11,126 samples, SNVs for 10,418 samples, and clinical data (including survival data) for all 10,005 samples. Supplementary Table 6 summarizes the data obtained from TCGA.

### Cancer morphological type and anatomical location

The cell type of origin and the anatomical site of cancer occurrence can reflect in similarities in tumor incidence and behaviour. In the past decade it has been shown for many different cancers that such similarities correlate with similar patterns of gene expression, epigenetics and characteristic chromosome abnormalities that link the cell of origin with the tumor [9]. We used information on the cell type of origin and anatomical site of tumor occurrence in building our model for predicting survival, integrating annotations for each tumor into the GCN as described below.

While it is difficult to unambiguously assign a cell of origin to all cancers, we can make use of the morphological characterization of tumors available in the NCI Thesaurus terminology which carries within it an implied association with cell type or tissue of origin [64]. For example, carcinoma (C2916) has as parent in the NCI Thesaurus “epithelial neoplasm” (C3709) which captures information on the cell type of origin. For each tumor, the most primitive parent below the superclass of “Neoplasm by morphology” or “Neoplasm by site” was used to describe the tumor sample. Tumors were annotated to 33 NCIT classes in the data provided by TGCA. The only exception was for tumors of neural crest origin where this was considered to be a more meaningful classification of these tumors given the close similarity between the ontogeny, behaviour and characteristics of these tumor types [65]. This concept is not available in either NCIT or in ICD-O3 but allowed us to express the similarity between for example melanoma whose parent class in the morphological axis is only “melanocytic neoplasm” and adrenal pheochromocytoma, classified only as an “epithelial neoplasm”.

Tumor topography presents a different set of problems in that it may be variously characterized as the site of origin of a specific instance of the tumor or the site of the originating cell. For example, using ICD-O3 [66], osteosarcoma of the kidney can be described as either located in the kidney or in bone [67]. The TGCA data was coded to ICD-O3 in the sense of the location of the tumor or site of biopsy. For the most part, these are primary tumors found in the tissue location of the presumed cell of origin. However, with lymphoid neoplasms arising in lymphoid tissues around the body, these are not annotated to the reticuloendothelial system or the blood, but the organ in which they were found. We have consequently used most of the 51 TGCA anatomical annotations as given, as they capture information from an orthogonal axis to the morphological characteristics, and it is known that site of occurrence often has a characteristic effect on tumor behavior. In only a few cases did we make changes: annotations to “Uterus NOS” and “Corpus Uteri” were merged, as were annotations of cholangiocarcinoma to “Liver and intrahepatic bile ducts” and “Other and unspecified parts of biliary tract”. Rectal adenocarcinoma tumors annotated to “connective, subcutaneous and other soft tissues” were reassigned to “Rectum”. Supplementary Tables 7 and 8 show the assignment of cancers to morphological type and anatomical parts.

### Protein–Protein interaction data

We use a protein interaction network for human proteins downloaded on 29 April 2020 from the STRING database version 11.0 [46]. STRING 11.0 contains 19,257 proteins and 11,780,842 edges between them incorporating both direct physical interactions and other functional interactions. STRING provides a confidence score for each interaction. We remove interactions with confidence score of less than 700. The remaining interaction network consists of 17,186 proteins with 736,125 interactions. We map the protein identifiers in the STRING interaction network to Ensembl gene identifiers [68] resulting in one gene for each protein. Nodes in our graph aim to represent a combination of genes, transcripts, and proteins.

### Cancer drivers and prognostic markers

We retrieved the driver genes for each cancer type from COSMIC database [69] on 13 June 2020. COSMIC contains a total of 723 driver genes within 327 cancer types. We mapped 359 driver genes to the 33 cancer types in TCGA.

We further used the Personal Cancer Genome Reporter (PCGR) [70] version 0.9.0 on 4 October 2020, which is a functional annotation tool to interpret somatic SNVs and CNVs. The PCGR tool combines several knowledge resources of tools and databases such as Variant Effect Predictor (VEP) [71], CHASMplus [72], Cancer Genome Interpreter database (CGI) [73] and TCGA which produce an individual specific report for all the 33 cancer types. We obtained a total of 135 prognostic markers including 69 driver genes and then derived their averaged rank using the attention mechanism as shown in Supplementary Table 5.

### Processing of multi-omics data

#### Absolute Gene Expression Data

TCGA provides gene expression data for cancer samples as read counts normalized by different approaches: Fragments per Kilobase of transcript per Million mapped reads (FPKM), and the upper quartile of Fragments per Kilobase of transcript per Million mapped reads (FPKM-UQ). FPKM normalizes read count by dividing it by the gene length and the total number of reads mapped to protein-coding genes. The FPKM-UQ is a modified FPKM calculation in which the total protein-coding read count is replaced by the 75th percentile read count value for the sample. We assigned the provided gene expression values for each sample to the gene entities by direct match (i.e., a 1-1 mapping) of Ensembl gene identifiers as provided in TCGA dataset and gene nodes in the patient graph. In total, we assigned all graph nodes (i.e., the 17,186 genes) to their expression values.

#### Differential Gene Expression Data

We applied differential expression analysis to identify differentially expressed genes using the TCGAbiolinks library [74], which calculates the difference of expression level of a gene between the mutant and normal sample multiplied by the *log2 Fold Change (log2FC)* between normal *A* and tumor *B* tissues for each sample:

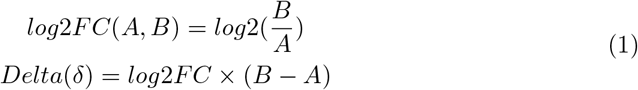

We assign the calculated *δ* values for all the 17,186 gene nodes.

#### Absolute DNA Methylation Data

Methylation is a biological process in which methyl groups are added to the DNA molecules by enzymes that affect (i.e., methylate) specific DNA regions (called CpG sites) which in turn change how genes being expressed and regulated [75]. TCGA provides a measurement for the level of methylation at known CpG sites as beta values, i.e., the ratio between the methylated probe intensity and the overall intensity (i.e., sum of methylated and unmethylated probe intensities) [76]. It falls between 0 (lower levels of methylation) and 1 (higher levels of methylation). We mapped the provided level of methylation values for transcript entities *t*_*i*_ in the TCGA dataset to their corresponding gene nodes in the patient graph by averaging the methylation values for these transcripts and assigned the resulted value to their corresponding gene as follow:

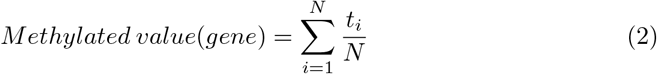

In total, 863,904 CpG loci have been sequenced and we assign all graph nodes (i.e., 17,186 genes) to their averaged transcripts methylated values.

#### Differential DNA Methylation Data

We identified differentially methylated regions (DMR) by measuring the significant difference between the methylated value in tumor and normal tissue. We consider regions as differentially methylated based on the Wilcoxon rank-sum test adjusted by Benjamini-Hochberg method with *p <* 0.05, and then we assign the calculated p-value for each gene nodes (i.e., 17,186 genes).

#### Copy Number Variation Data

The TCGA dataset provides categorical data for each gene whether the gene is in a copy number gain (value of −1), loss (value of 1), or not (value of 0). We mapped the provided CNV category for each gene in the TCGA dataset to the gene nodes in the patient graph (i.e., 17,186 genes).

#### Single Nucleotide Variation Data

TCGA provides single nucleotide variants (SNVs) for germline and somatic variants. We annotated each variant with its pathogenicity score derived from the FATHMM tool [77] using Annovar [78]. We then assigned each gene node with the maximum pathogenicity score among all its variants, separately for germline and somatic variants; if a variant is intergenic, we assign its pathogenicity score to the nearest gene. The pathogenicity values range from 0 to 1.

#### Clinical Data

Clinical data provided by TCGA includes several types of data such as patient diagnosis, demographics, exposures, laboratory tests, and family relationships, age, survival time, and the number of days to last followup. We use the days to last followup and days to death as patients survival time values assigned to each patient graph whether this patient alive or dead.

### Mapping sample features to node features

We assign values derived from different omics data types to the STRING graph. Each TCGA sample is used to assign a set of attributes to nodes in the graph. We define a set of mappings functions *f*_*i*_ : *S* ↦*G* that map information derived from an individual sample *S* to attributes of nodes in *G*. We implement mapping functions for gene expression, methylation, somatic mutations, and copy number variants.

We normalize the data when mapping them to our graph; here, normalization means to transform the values so they lie in a range between 0 and 1. There are different ways in which we can perform this normalization: globally, by gene or node, and by sample. A global normalization identifies the minimum and maximum values *ν*_*min*_ and *ν*_*max*_ of gene expression across all samples and all genes and normalizes all values based on *ν*_*min*_ and *ν*_*max*_. Normalization by gene identifies the minimum and maximum expression values 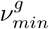 and 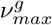 for a gene *g* across all samples, and normalizes the expression values for each gene *g* based on 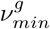 and 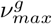. Normalization by sample identifies the minimum and maximum values *ν*_*min*_ and *ν*_*max*_ of gene expression across all genes for that sample and normalizes all values based on *ν*_*min*_ and *ν*_*max*_.

Gene-based normalization captures the range of gene expression (or methylation) across multiple samples and can be used to determine whether a gene is expressed relatively high or low in a single sample compared to other samples. Sample-based normalization, on the other hand, identifies the minimum and maximum gene expression within each sample and normalizes each expression within a sample by these values; sample-based normalization can be used to determine genes expressed relatively high or low within a sample compared to other genes in the same sample. Each of these normalization techniques alone is subject to several biases, and we can combine the different normalization methods and assign multiple attributes to each node in *G*.

### The model architecture

Our model combines a Cox proportional hazards regression with a graph convolutional network that incorporates prior knowledge as shown in Figure 1. The model takes as input omics data derived from individual cancer samples (gene expression, DNA methylation, CNVs, SNVs), the general type of cancer taken (one of the 33 types in TCGA), the anatomical location of the cancer sample, the cancer subtype (i.e., the cancer subgroup based on certain characteristics of the cancer cells), and the cell type of origin (describe from which cell this cancer originate). For using the model, only the omics data, cancer type, and anatomical location must be provided whereas the morphological classification is derived automatically according to Supplementary Table 7. As an output, our model produces a prediction of survival time for a patient based on the chosen different cancer types and subtypes.

#### Graph Convolutional Network

We use a Graph Convolutional Network (GCN) [79] to process the omics data. A GCN is a neural network that operates on graphs. A GCN uses as inputs a graph *G* = (*V, E*) and a feature matrix *X* of dimension |*V* |× |*F* | (where |*V*| is the number of nodes in *G* and |*F* |is the number of features per node). The matrix *A* of dimension |*V* |× |*V*| is the adjacency matrix of *G*.

In our model, patient-derived omics data is represented as a feature matrix *X* of the form 17186 ×*ζ* where 17186 is the number of nodes in our graph and *ζ* the number of features we assign for each node; depending on the model and availability of data for one sample, we assign between 1 and 8 features to each node. An adjacency matrix of the form 17186× 17186 is used to represent the graph.

The adjacency and feature matrices are used as input to a GCN layer, *H*^1^ = *f* (*X, A*) with *f* being a propagation rule; we use *f* as 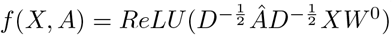, where *RelU* (*x*) = max(0, *x*) is the activation, Â = *A* + *I* is the adjacency matrix with inserted self-loops, *D*_*ii*_ = Σ_*j*=0_ Â_*ij*_ is the degree matrix, and *W* ^0^ is the weight matrix for the first layer.

Following the first graph convolutional layer, we apply a pooling operation based on self attention [80]. The self attention score *Z* is calculated as

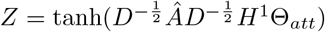, *idx* = *topRank*(*Z*; ⌈*k* · *V* ⌉); *Z*_*mask*_ = *Z*_*idx*_ with 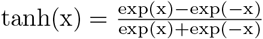 is the activation function, 0 *< k* ≤1, *k* being the pooling ratio that determines the number of nodes to maintain, top ⌈*k* · *V* ⌉ nodes are selected based on the value of *Z, topRank* is a function that return the indices of the top ⌈*k* · *V* ⌉ nodes, *Z*_*mask*_ is the mask for feature attention, and *idx* is an indexing operation. The resulted attention score matrix is of dimension *V* ×1.

We apply second graph convolutional layer on the pooled graph. The input for the second GCN are the adjacency and feature matrices for the pooled graph, *H*^2^ = *f* (*X*_*p*_, *A*_*p*_); using the propagation rule *f* here as 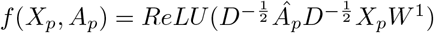.

Then, we add a fully connected layer that applies a linear transformation on the matrix *X* after the convolution as *Y* = *W*^*l*^*X* + *b*, where *W*^*l*^ is the weight for the fully connected layer, and *b* is the learnable bias. Subsequently, we apply a sigmoid function that transforms *Y* to be between 0 and 1.

### Cox regression

Survival prediction involves censored data where either an event is observed in a particular time or no event is observed. The Cox regression model is semi-parametric as there is no assumption about the distribution of the outcome. For a given patient *i* at time of an event *t* (either death or censored), the hazard function *h*(*t, X*_*i*_) in the Cox model is build upon the proportional hazards assumption expressed as:

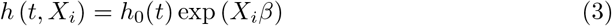

where *i* = 1, 2, …, *N, h*_0_(*t*) is the baseline hazard function, *X*_*i*_ = (*x*_*i*1_, *x*_*i*2_, · · ·, *x*_*iN*_) are corresponding to the covariates for an individual sample and *β* = (*β*_1_, *β*_2_,, *β*_*N*_) are the model coefficients.

### Tuning the model

We utilized the RayTune tool [81] for choosing optimal sets of hyperparameters of the graph convolutional network, we tuned the number of layers (1-5 layers), the respective kernel sizes *K*∈ {4, 8, 16, 32, 64}and the number of self-attention pooling layers *P* ∈ {1, 2, 3, 4}. The optimal parameters were: 2-convolutional layers, kernel of size 32 for the first layer and 16 for the second layer, and one self-attention pooling layer.

Furthermore, we investigate different graph-based architectures such as GENConv [82], GraphSAGE [83], GraphConv [84] and APPNP [85]. In GENConv, a deeper GCN architecture is used with the help of residual connections. Furthermore, GENConv propose a generalized message aggregation function which relied on permutation invariant functions. For this architecture, we tuned the number of layers (1 − 5 layers) and the respective kernel sizes *KN* ∈ {4, 8, 16, 32, 64} and the aggregation schema *AG* ∈{*softmax, softmax*_*s*_*g, power, add, mean, max*}. The optimal set of parameters were: 2-convolutional layers with size of kernel equal 64 in the first layer and 32 in the second layer and using *mean* as an aggregation operation. In GraphSAGE, they introduced an inductive classification task where the goal is to generalize the graph information to unseen nodes interactions during training. For this architecture, we tuned the number of layers (1-5 layers) and the respective kernel sizes *KN* ∈{4, 8, 16, 32, 64}. The optimal set of parameters were: 2-convolutional layers with size of kernel equal 16 in the first layer and 8 in the second layer. In GraphConv, they introduced a hierarchical way (i.e., Weisfeiler-Lehman (WL) graph isomorphism test) to generalize message passing process to higher orders of learn features for sub-graphs than vertices. For this architecture, we tuned the number of layers (1-5 layers), the respective kernel sizes *KN* ∈{4, 8, 16, 32, 64}and the aggregation schema *AG* ∈ {*add, mean, max*}. The optimal set of parameters were: 1-layer of kernel size 16 and using *max* as an aggregation operation. In APPNP, the graph convolutions are defined with a teleport probability *α* inspired by the original PageRank algorithm [86]. For this architecture, we tuned the number of layers (1-5 layers), the respective kernel sizes *KN*∈ {4, 8, 16, 32, 64}, the propagation steps *K*∈ {1, 2, ……, 10}and the teleport probability *α*∈ (0, 1] as used to perform tuning in APPNP paper. The optimal set of parameters were: 1-layer, kernel size of 32, propagation steps of 3 and teleport probability of 0.2. Supplementary Table 9 summarized the evaluation results between different graph-based architectures.

### Training, validation and testing

We investigated the performance of our deep learning-based regression algorithm in predicting survival probability for a patient being survived. In our experiments, we used samples omics data within 33 different cancer types with their known survival time which defined as either the days until the patient’s death or until their last follow-up. Both input data (i.e., 4 types of omics data) and output (i.e., the probability of whether a patient survived or not) were standardized to mean of zero and standard deviation of one. We randomly split our datasets into 85% and 15%, respectively, and we used 15% of the training set as a validation set. The training and validation sets are used to train and tune model parameters and select the best models, while the test set has been used for reporting the evaluation results. We implemented our model using PyTorch Geometric (PyG) [87] and Pycox [88] and performed training on Nvidia Tesla V100 GPUs which takes 1.5 hours. We utilized RayTune for tuning models parameters (see Tuning the model subsection). We used Adam to optimize the graph convolutional network parameters in training, and to predict the survival probability for a patient, we train the graph network as a regression task using partial negative likelihood:

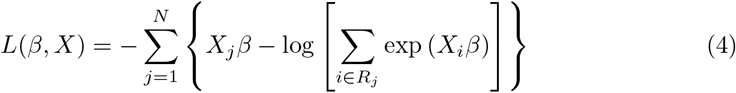

where *i* = 1, 2, …, *N, X*_*i*_ are corresponding to the covariates for an individual sample, *β* are the Cox model parameters, *U* is the set of uncensored samples, and *R*_*i*_ is the set of patients with survival times *Y*_*j*_ ⩾*Y*_*i*_.

For the evaluation of our model and other different tested models, we use the Concordance index (C-index) [89] as shown in Equation 5 which measures the concordance between actual survival time and predicted hazard scores of all pairs of individuals. C-index is an appropriate measurement in capturing the discriminating ability of a predictive covariate to separate individuals with longer survival from those with shorter survival when predicting their survival time [90]. In addition, we use the Root Mean Square Error (RMSE) which measures the square root of the average difference between the predicted hazard scores values and the actual survival time values. The C-index is computed as

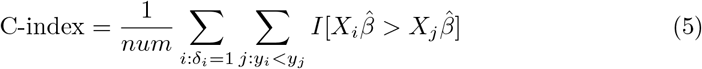

where *i, j* ∈ 1, …, *N, num* denotes the number of all comparable pairs, *I*[·] is the indicator function and 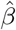 is the estimated parameters from the Cox based models.

Furthermore, we perform a random assignment for the omics features (i.e., assign randomly the features to different samples than the original one). The aim of doing this experiment is to test how our model will perform compared to the correct assignment, and whether it predicts based on spurious correlations introduced through the graph.

We find (Supplementary Table 1) that the prediction performance by applying random assignment is significantly different (lower) both for the C-index and RMSE results than the original assignment in the three tested cancer types (Breast data, RMSE: *p* = 0.0408 C-index: *p* = 0.0441, Lung data, RMSE: *p* = 0.0438 C-index: *p* = 0.0426, Glioblastoma data, RMSE: *p* = 0.0401 C-index: *p* = 0.0421, two-tailed t-test).

### Analysis of similarities and attention ranking

To estimate the similarity between representations generated for different cancer types, we compute Pearson correlation among the element-wise arithmetic mean of the representations generated from each sample.

## Availability of data and software

All data and software used to develop, apply and evaluate the models, except data obtained from TCGA, are freely available at

https://github.com/bio-ontology-research-group/DeepMOCCA. Omics data used to generate and apply the models is available from The Cancer Genome Atlas data portal for researchers which have approved access by the NCI Data Access Committee.

## Ethical approval

This work has been reviewed and approved by the Institutional Bioethics Committee at King Abdullah University of Science and Technology on 31 January 2019 under approval number 19IBEC02. Access to genomic data from The Cancer Genome Atlas was approved by the NCI Data Access Committee under Project ID 18502 “Machine learning for prioritization of causal variants in Mendelian and oligogenic disease”.

## Acknowledgements

We acknowledge the use of computational resources from the KAUST Supercomputing Core Laboratory.

The results published here are in whole or part based upon data generated by The Cancer Genome Atlas managed by the NCI and NHGRI. Information about TCGA can be found at http://cancergenome.nih.gov.

