## Supplementary Tables for "DeepMOCCA: A pan-cancer prognostic model identifies personalized prognostic markers through graph attention and multi-omics data integration"

[2]College of Medical and Dental Sciences, Institute of Cancer and Genomic Sciences, University of Birmingham, B15 2TT, Birmingham, United Kingdom, and Institute of Translational Medicine, University Hospitals Birmingham, NHS Foundation Trust, B15 2TT, Birmingham, United Kingdom, and NIHR Biomedical Research Centre, B15 2TT, Birmingham, United Kingdom, and NIHR Experimental Cancer Medicine Centre, B15 2TT, Birmingham, United Kingdom, and MRC Health Data Research UK (HDR UK) Midlands, B15 2TT, Birmingham, United Kingdom

[3]Department of Physiology, Development, and Neuroscience, University of Cambridge, Downing Street, CB2 3EG, Cambridge, UK

[*]

| | | | Breast | | Lung | | Glioblastoma | |
|---|---|---|---|---|---|---|---|---|
| | | Number of samples | 1044 | | 509 | | 166 | |
| | | Type of normalization | Min-max normalization | Log normalization | Min-max normalization | Log normalization | Min-max normalization | Log normalization |
| Absolute gene expression (FPKM-UQ — Column+Row) | Original assignment | RMSE | 0.1623 ± 0.0263 | 0.0792 ± 0.0063 | 0.0687 ± 0.0047 | 0.0454± 0.0021 | 0.1738 ± 0.0302 | 0.1504 ± 0.0226 |
| | | C-index | 0.5943 | 0.6634 | 0.5920 | 0.6298 | 0.5734 | 0.6077 |
| | Random assignment | RMSE | 0.1980 ± 0.0392 | 0.1517 ± 0.0230 | 0.4954 ± 0.2454 | 0.4327 ± 0.1872 | 0.5852 ± 0.3425 | 0.3548 ± 0.1259 |
| | | C-index | 0.5963 | 0.5905 | 0.5874 | 0.5982 | 0.5109 | 0.5645 |
| Differential gene expression (FPKM-UQ — Column+Row) | Original assignment | RMSE | 0.1537 ± 0.0236 | 0.0599 ± 0.0036 | 0.0544 ± 0.0030 | 0.0328 ± 0.0011 | 0.1386 ± 0.0192 | 0.1191 ± 0.0142 |
| | | C-index | 0.6358 | 0.6692 | 0.6226 | 0.6487 | 0.5854 | 0.5990 |
| | Random assignment | RMSE | 0.0834 ± 0.0070 | 0.1843 ± 0.0340 | 0.1274 ± 0.0162 | 0.2089 ± 0.0436 | 0.3460 ± 0.1197 | 0.3079 ± 0.0948 |
| | | C-index | 0.5837 | 0.5964 | 0.6028 | 0.5647 | 0.5083 | 0.5274 |
| Absolute gene expression + Differential gene expression | Original assignment | RMSE | 0.0588 ± 0.0035 | 0.0375 ± 0.0014 | 0.0468 ± 0.0022 | 0.0385 ± 0.0015 | 0.1037 ± 0.0108 | 0.0920 ± 0.0085 |
| | | C-index | 0.6717 | 0.7053 | 0.6537 | 0.6764 | 0.6036 | 0.6215 |
| | Random assignment | RMSE | 0.1265 ± 0.0160 | 0.2761 ± 0.0762 | 0.2475 ± 0.0613 | 0.3572 ± 0.1276 | 0.3378 ± 0.1141 | 0.2847 ± 0.0811 |
| | | C-index | 0.5632 | 0.6196 | 0.6026 | 0.6275 | 0.5357 | 0.5380 |
| Absolute DNA methylation (FPKM-UQ — Column+Row) | Original assignment | RMSE | 0.0639 ± 0.0041 | 0.0567 ± 0.0032 | 0.0347 ± 0.0012 | 0.0290 ± 0.0008 | 0.0927 ± 0.0086 | 0.0753 ± 0.0057 |
| | | C-index | 0.6836 | 0.7087 | 0.6756 | 0.6924 | 0.6443 | 0.6564 |
| | Random assignment | RMSE | 0.0993 ± 0.0099 | 0.2354 ± 0.0554 | 0.2832 ± 0.0802 | 0.2538 ± 0.0644 | 0.5952 ± 0.3543 | 0.5511 ± 0.3037 |
| | | C-index | 0.5124 | 0.6328 | 0.6042 | 0.6268 | 0.5365 | 0.5376 |
| Differential DNA methylation (FPKM-UQ — Column+Row) | Original assignment | RMSE | 0.0491 ± 0.0024 | 0.0265 ± 0.0007 | 0.0304 ± 0.0009 | 0.0293 ± 0.0009 | 0.1038 ± 0.0108 | 0.0821 ± 0.0067 |
| | | C-index | 0.7055 | 0.7236 | 0.6826 | 0.6972 | 0.6384 | 0.6608 |
| | Random assignment | RMSE | 0.6410 ± 0.4109 | 0.4792 ± 0.2296 | 0.2281 ± 0.0520 | 0.3168 ± 0.1004 | 0.4653 ± 0.2165 | 0.4239 ± 0.1797 |
| | | C-index | 0.6052 | 0.6174 | 0.6297 | 0.5727 | 0.5837 | 0.5913 |
| Absolute DNA methylation + Differential DNA methylation | Original assignment | RMSE | 0.0364 ± 0.0013 | 0.0319 ± 0.0010 | 0.0264 ± 0.0007 | 0.0223 ± 0.0005 | 0.0684 ± 0.0047 | 0.0545 ± 0.0030 |
| | | C-index | 0.7374 | 0.7405 | 0.7043 | 0.7169 | 0.6662 | 0.6775 |
| | Random assignment | RMSE | 0.3137 ± 0.0984 | 0.4248 ± 0.1805 | 0.2638 ± 0.0696 | 0.3587 ± 0.1287 | 0.4032 ± 0.1626 | 0.3892 ± 0.1515 |
| | | C-index | 0.6321 | 0.5994 | 0.6104 | 0.6273 | 0.5527 | 0.5786 |
| CNV | Original assignment | RMSE | 0.2738 ± 0.0750 | 0.3053 ± 0.0932 | 0.1994 ± 0.0398 | 0.1736 ± 0.0301 | 0.4735 ± 0.2242 | 0.3806 ± 0.1449 |
| | | C-index | 0.5473 | 0.5547 | 0.5723 | 0.6044 | 0.5276 | 0.5416 |
| | Random assignment | RMSE | 0.7784 ± 0.6059 | 0.5638 ± 0.3179 | 0.5487 ± 0.3011 | 0.4885 ± 0.2386 | 0.7950 ± 0.6320 | 0.7358 ± 0.5414 |
| | | C-index | 0.5143 | 0.6058 | 0.5322 | 0.5136 | 0.5027 | 0.5144 |
| VCF (combine germline + somatic mutations separately) | Original assignment | RMSE | 0.3754 ± 0.1409 | 0.2962 ± 0.0877 | 0.2183 ± 0.0477 | 0.1829 ± 0.0335 | 0.2875 ± 0.0827 | 0.2526 ± 0.0638 |
| | | C-index | 0.5439 | 0.5544 | 0.5679 | 0.5815 | 0.5462 | 0.5732 |
| | Random assignment | RMSE | 0.4068 ± 0.1655 | 0.3472 ± 0.1205 | 0.6366 ± 0.4053 | 0.6018 ± 0.3622 | 0.7159 ± 0.5125 | 0.7006 ± 0.4908 |
| | | C-index | 0.5276 | 0.5584 | 0.5034 | 0.5366 | 0.5028 | 0.5169 |
| Differential gene expression + Differential DNA methylation | Original assignment | RMSE | 0.0268 ± 0.0007 | 0.0189 ± 0.0004 | 0.0245 ± 0.0006 | 0.0187 ± 0.0003 | 0.0304 ± 0.0023 | 0.2257 ± 0.0509 |
| | | C-index | 0.7784 | 0.7902 | 0.7913 | 0.8054 | 0.7065 | 0.7251 |
| | Random assignment | RMSE | 0.0753 ± 0.0057 | 0.0943 ± 0.0089 | 0.0820 ± 0.0067 | 0.0632 ± 0.0040 | 0.2482 ± 0.0616 | 0.1809 ± 0.0327 |
| | | C-index | 0.6269 | 0.6411 | 0.6407 | 0.6712 | 0.5561 | 0.5856 |
| Differential gene expression + Differential DNA methylation+CNV | Original assignment | RMSE | 0.0148 ± 0.0002 | 0.0106 ± 0.0001 | 0.0141 ± 0.0002 | 0.0138 ± 0.0002 | 0.0285 ± 0.0008 | 0.0173 ± 0.0003 |
| | | C-index | 0.8330 | 0.8529 | 0.8496 | 0.8526 | 0.7662 | 0.7916 |
| | Random assignment | RMSE | 0.4820 ± 0.2323 | 0.5372 ± 0.2886 | 0.1962 ± 0.0385 | 0.1651 ± 0.0273 | 0.0824 ± 0.0068 | 0.0754 ± 0.0057 |
| | | C-index | 0.6528 | 0.6185 | 0.6289 | 0.6572 | 0.5823 | 0.6107 |
| Differential gene expression + Differential DNA methylation+CNV+VCF (separately) | Original assignment | RMSE | 0.0103 ± 0.0001 | **0.0101 ± 0.0001** | 0.0115 ± 0.0001 | **0.0111 ± 0.0001** | 0.0219 ± 0.0005 | **0.0159 ± 0.0002** |
| | | C-index | 0.8378 | **0.8593** | 0.8582 | **0.8663** | 0.7804 | **0.7941** |
| | Random assignment | RMSE | 0.6747 ± 0.4552 | 0.4733 ± 0.2240 | 0.1126 ± 0.0127 | 0.1106 ± 0.0123 | 0.0953 ± 0.0091 | 0.0877 ± 0.0077 |
| | | C-index | 0.5378 | 0.5947 | 0.5647 | 0.5870 | 0.6042 | 0.6161 |
| | | RMSE p-value | 0.0424 | 0.0408 | 0.0480 | 0.0438 | 0.0417 | 0.0401 |
| | | C-index p-value | 0.0453 | 0.0441 | 0.0434 | 0.0426 | 0.0445 | 0.0421 |

Table 1: Performance comparison on different combination of features trained individually on Breast, Lung and Glioblastoma

| | | | | Breast | | Lung | | Glioblastoma | |
|---|---|---|---|---|---|---|---|---|---|
| | | | Number of samples | 1044 | | 509 | | 166 | |
| | | | Type of normalization | Min-max normalization | Log normalization | Min-max normalization | Log normalization | Min-max normalization | Log normalization |
| Absolute gene expression | Matrix | Read Count | RMSE | 0.3932 ± 0.1546 | 0.2735 ± 0.0748 | 0.5397 ± 0.2913 | 0.4734 ± 0.2241 | 0.7374 ± 0.5438 | 0.6369 ± 0.4056 |
| | | | C-index | 0.5384 | 0.5963 | 0.5284 | 0.5526 | 0.5159 | 0.5454 |
| | | FPKM | RMSE | 0.2433 ± 0.0592 | 0.2174 ± 0.0473 | 0.2734 ± 0.0747 | 0.2285 ± 0.0522 | 0.6826 ± 0.4654 | 0.5315 ± 0.2825 |
| | | | C-index | 0.5573 | 0.6184 | 0.5447 | 0.5578 | 0.5288 | 0.5598 |
| | | FPKM-UQ | RMSE | 0.1572 ± 0.0247 | 0.2045 ± 0.0418 | 0.1435 ± 0.0206 | 0.1285 ± 0.0165 | 0.5290 ± 0.2798 | 0.4360 ± 0.1901 |
| | | | C-index | 0.5792 | 0.6397 | 0.5732 | 0.5954 | 0.5416 | 0.5723 |
| | Column | Read Count | RMSE | 0.2210 ± 0.0488 | 0.2167 ± 0.0470 | 0.3651 ± 0.1333 | 0.2592 ± 0.0672 | 0.5360 ± 0.2873 | 0.3862 ± 0.1492 |
| | | | C-index | 0.5547 | 0.6282 | 0.5383 | 0.5626 | 0.5277 | 0.5589 |
| | | FPKM | RMSE | 0.1745 ± 0.0305 | 0.1164 ± 0.0135 | 0.1908 ± 0.0364 | 0.1258 ± 0.0158 | 0.2758 ± 0.0761 | 0.1673 ± 0.0280 |
| | | | C-index | 0.5673 | 0.6301 | 0.5695 | 0.5931 | 0.5439 | 0.5728 |
| | | FPKM-UQ | RMSE | 0.1587 ± 0.0252 | 0.0863 ± 0.0074 | 0.0962 ± 0.0093 | 0.0523 ± 0.0027 | 0.1472 ± 0.0217 | 0.0945 ± 0.0089 |
| | | | C-index | 0.5934 | 0.6567 | 0.5820 | 0.6148 | 0.5724 | 0.6004 |
| | Row | Read Count | RMSE | 0.3121 ± 0.0974 | 0.2592 ± 0.0672 | 0.2615 ± 0.0684 | 0.1945 ± 0.0378 | 0.6972 ± 0.4861 | 0.5320 ± 0.2830 |
| | | | C-index | 0.5391 | 0.6037 | 0.5248 | 0.5494 | 0.5151 | 0.5482 |
| | | FPKM | RMSE | 0.1862 ± 0.0347 | 0.1124 ± 0.0126 | 0.1872 ± 0.0350 | 0.1383 ± 0.0191 | 0.5663 ± 0.3207 | 0.3857 ± 0.1488 |
| | | | C-index | 0.5638 | 0.6244 | 0.5586 | 0.5729 | 0.5377 | 0.5535 |
| | | FPKM-UQ | RMSE | 0.1676 ± 0.0281 | 0.1097 ± 0.0120 | 0.1621 ± 0.0263 | 0.1278 ± 0.0163 | 0.1706 ± 0.0291 | 0.1446 ± 0.0209 |
| | | | C-index | 0.5720 | 0.6486 | 0.5728 | 0.5853 | 0.5629 | 0.6062 |
| | Column+Row | Read Count | RMSE | 0.1872 ± 0.0350 | 0.1933 ± 0.0374 | 0.2063 ± 0.0426 | 0.1646 ± 0.0271 | 0.5421 ± 0.2939 | 0.5739 ± 0.3294 |
| | | | C-index | 0.5783 | 0.6491 | 0.5540 | 0.5729 | 0.5450 | 0.5656 |
| | | FPKM | RMSE | 0.1880 ± 0.0353 | 0.1081 ± 0.0117 | 0.1105 ± 0.0122 | 0.0925 ± 0.0086 | 0.2218 ± 0.0492 | 0.1572 ± 0.0247 |
| | | | C-index | 0.5848 | 0.6424 | 0.5818 | 0.6188 | 0.5510 | 0.5791 |
| | | FPKM-UQ | RMSE | 0.1623 ± 0.0263 | **0.0792 ± 0.0063** | 0.0687 ± 0.0047 | **0.0454± 0.0021** | 0.1738 ± 0.0302 | **0.1504 ± 0.0226** |
| | | | C-index | 0.5943 | **0.6634** | 0.5920 | **0.6298** | 0.5734 | **0.6077** |
| | Matrix+Column+Row | Read Count | RMSE | 0.2054 ± 0.0422 | 0.2358 ± 0.0556 | 0.3775 ± 0.1425 | 0.2874 ± 0.0826 | 0.4958 ± 0.2458 | 0.3253 ± 0.1058 |
| | | | C-index | 0.5563 | 0.6061 | 0.5339 | 0.5687 | 0.5225 | 0.5598 |
| | | FPKM | RMSE | 0.0944 ± 0.0089 | 0.1937 ± 0.0375 | 0.1706 ± 0.0291 | 0.1159 ± 0.0134 | 0.5691 ± 0.3239 | 0.4873 ± 0.2375 |
| | | | C-index | 0.5619 | 0.6236 | 0.5554 | 0.5821 | 0.5327 | 0.5668 |
| | | FPKM-UQ | RMSE | 0.0930 ± 0.0086 | 0.1653 ± 0.0373 | 0.1361 ± 0.0185 | 0.0843 ± 0.0071 | 0.1439 ± 0.0207 | 0.1146 ± 0.0131 |
| | | | C-index | 0.5837 | 0.6527 | 0.5748 | 0.6106 | 0.5760 | 0.5915 |
| Differential gene expression | Matrix | Read Count | RMSE | 0.3468 ± 0.1203 | 0.2784 ± 0.0775 | 0.4862 ± 0.2364 | 0.3498 ± 0.1224 | 0.7528 ± 0.5667 | 0.5674 ± 0.3219 |
| | | | C-index | 0.5416 | 0.5925 | 0.5446 | 0.5703 | 0.5271 | 0.5638 |
| | | FPKM | RMSE | 0.2168 ± 0.0470 | 0.1843 ± 0.0340 | 0.2584 ± 0.0668 | 0.1952 ± 0.0381 | 0.6309 ± 0.3980 | 0.4622 ± 0.2136 |
| | | | C-index | 0.5704 | 0.6319 | 0.5539 | 0.5729 | 0.5567 | 0.5646 |
| | | FPKM-UQ | RMSE | 0.1793 ± 0.0321 | 0.1697 ± 0.0288 | 0.1392 ± 0.0194 | 0.1044 ± 0.0109 | 0.5282 ± 0.2790 | 0.3899 ± 0.1520 |
| | | | C-index | 0.5774 | 0.6472 | 0.5776 | 0.6274 | 0.5478 | 0.5824 |
| | Column | Read Count | RMSE | 0.2074 ± 0.0430 | 0.1783 ± 0.0318 | 0.3298 ± 0.1088 | 0.2453 ± 0.0602 | 0.5077 ± 0.2578 | 0.3347 ± 0.1120 |
| | | | C-index | 0.5692 | 0.6408 | 0.5529 | 0.5926 | 0.50264 | 0.5881 |
| | | FPKM | RMSE | 0.1756 ± 0.0308 | 0.0965 ± 0.0093 | 0.1385 ± 0.0192 | 0.1163 ± 0.0135 | 0.2586 ± 0.0669 | 0.1396 ± 0.0195 |
| | | | C-index | 0.6063 | 0.6592 | 0.5942 | 0.6184 | 0.5647 | 0.6068 |
| | | FPKM-UQ | RMSE | 0.6371 ± 0.4059 | 0.1533 ± 0.0235 | 0.0427 ± 0.0018 | 0.0361 ± 0.0013 | 0.1173 ± 0.0138 | 0.0734 ± 0.0054 |
| | | | C-index | 0.6135 | 0.6453 | 0.6256 | 0.6421 | 0.5920 | 0.6647 |
| | Row | Read Count | RMSE | 0.2957 ± 0.0874 | 0.2288 ± 0.0523 | 0.2163 ± 0.0468 | 0.1846 ± 0.0341 | 0.5830 ± 0.3399 | 0.2710 ± 0.0734 |
| | | | C-index | 0.5410 | 0.6165 | 0.5395 | 0.5582 | 0.5606 | 0.5705 |
| | | FPKM | RMSE | 0.1942 ± 0.0377 | 0.0895 ± 0.0080 | 0.1475 ± 0.0218 | 0.1372 ± 0.0188 | 0.5263 ± 0.2770 | 0.3632 ± 0.1319 |
| | | | C-index | 0.5787 | 0.6374 | 0.5814 | 0.6004 | 0.5651 | 0.5828 |
| | | FPKM-UQ | RMSE | 0.1780 ± 0.0317 | 0.0884 ± 0.0078 | 0.1195 ± 0.0143 | 0.1057 ± 0.0112 | 0.1339 ± 0.0179 | 0.1239 ± 0.0154 |
| | | | C-index | 0.5902 | 0.6559 | 0.6192 | 0.6237 | 0.5838 | 0.6147 |
| | Column+Row | Read Count | RMSE | 0.1873 ± 0.0351 | 0.0643 ± 0.0041 | 0.1705 ± 0.0291 | 0.1511 ± 0.0228 | 0.1818 ± 0.0331 | 0.1672 ± 0.0280 |
| | | | C-index | 0.6137 | 0.6618 | 0.5628 | 0.5825 | 0.5746 | 0.6054 |
| | | FPKM | RMSE | 0.1582 ± 0.0250 | 0.0748 ± 0.0056 | 0.0826 ± 0.0068 | 0.0572 ± 0.0033 | 0.1867 ± 0.0349 | 0.1017 ± 0.0103 |
| | | | C-index | 0.6024 | 0.6528 | 0.6246 | 0.6378 | 0.5758 | 0.5873 |
| | | FPKM-UQ | RMSE | 0.1537 ± 0.0236 | **0.0599 ± 0.0036** | 0.0544 ± 0.0030 | **0.0328 ± 0.0011** | 0.1386 ± 0.0192 | **0.1191 ± 0.0142** |
| | | | C-index | 0.6358 | **0.6692** | 0.6226 | **0.6487** | 0.5854 | **0.5990** |
| | Matrix+Column+Row | Read Count | RMSE | 0.2073 ± 0.0430 | 0.1953 ± 0.0381 | 0.2651 ± 0.0703 | 0.2644 ± 0.0699 | 0.4506 ± 0.2030 | 0.2613 ± 0.0683 |
| | | | C-index | 0.5749 | 0.6361 | 0.5474 | 0.5948 | 0.5365 | 0.5830 |
| | | FPKM | RMSE | 0.1946 ± 0.0379 | 0.0826 ± 0.0068 | 0.1643 ± 0.0270 | 0.1037 ± 0.0108 | 0.5272 ± 0.2779 | 0.4409 ± 0.1944 |
| | | | C-index | 0.5782 | 0.6306 | 0.5593 | 0.6085 | 0.5594 | 0.5772 |
| | | FPKM-UQ | RMSE | 0.1698 ± 0.0288 | 0.0622 ± 0.0039 | 0.1059 ± 0.0112 | 0.0415 ± 0.0017 | 0.1187 ± 0.0140 | 0.0935 ± 0.0087 |
| | | | C-index | 0.6432 | 0.6552 | 0.6266 | 0.6436 | 0.6133 | 0.6435 |

Table 2: Comparison of predictive performance using different normalization techniques on gene expression features trained individually on Breast, Lung and Glioblastoma.

| | | Morphological classification + Anatomical locations | |
|---|---|---|---|
| | Number of samples | RMSE | C-index |
| TCGA-ACC | 80 | 0.1163 ± 0.0135 | 0.7338 |
| TCGA-BLCA | 407 | 0.0947 ± 0.0090 | 0.7692 |
| TCGA-BRCA | 1044 | 0.0100 ± 0.0001 | 0.8597 |
| TCGA-CESC | 294 | 0.0847 ± 0.0072 | 0.8380 |
| TCGA-CHOL | 36 | 0.2864 ± 0.0820 | 0.6849 |
| TCGA-COAD | 433 | 0.0582 ± 0.0034 | 0.8661 |
| TCGA-DLBC | 37 | 0.1620 ± 0.0262 | 0.7405 |
| TCGA-ESCA | 184 | 0.1208 ± 0.0146 | 0.7847 |
| TCGA-GBM | 166 | 0.0122 ± 0.0001 | 0.8373 |
| TCGA-HNSC | 510 | 0.0114 ± 0.0001 | 0.8732 |
| TCGA-KICH | 66 | 0.1169 ± 0.0137 | 0.7728 |
| TCGA-KIRC | 339 | 0.0190 ± 0.0004 | 0.8676 |
| TCGA-KIRP | 288 | 0.0512 ± 0.0026 | 0.8023 |
| TCGA-LAML | 140 | 0.3178 ± 0.1010 | 0.7017 |
| TCGA-LGG | 511 | 0.1481 ± 0.0219 | 0.7652 |
| TCGA-LIHC | 371 | 0.1137 ± 0.0129 | 0.7303 |
| TCGA-LUAD | 509 | 0.0103 ± 0.0001 | 0.8781 |
| TCGA-LUSC | 496 | 0.0192 ± 0.0004 | 0.8621 |
| TCGA-MESO | 83 | 0.1346 ± 0.0181 | 0.7445 |
| TCGA-OV | 443 | 0.0163 ± 0.0003 | 0.8510 |
| TCGA-PAAD | 178 | 0.1675 ± 0.0281 | 0.7173 |
| TCGA-PCPG | 179 | 0.0287 ± 0.0008 | 0.8647 |
| TCGA-PRAD | 498 | 0.1842 ± 0.0339 | 0.7206 |
| TCGA-READ | 158 | 0.1480 ± 0.0219 | 0.7378 |
| TCGA-SARC | 255 | 0.0112 ± 0.0001 | 0.8735 |
| TCGA-SKCM | 456 | 0.2163 ± 0.0468 | 0.7336 |
| TCGA-STAD | 414 | 0.0273 ± 0.0007 | 0.8471 |
| TCGA-TGCT | 134 | 0.1376 ± 0.0189 | 0.6912 |
| TCGA-THCA | 496 | 0.0139 ± 0.0002 | 0.8483 |
| TCGA-THYM | 123 | 0.1277 ± 0.0163 | 0.6937 |
| TCGA-UCEC | 542 | 0.0925 ± 0.0086 | 0.7866 |
| TCGA-UCS | 55 | 0.1732 ± 0.0300 | 0.6608 |
| TCGA-UVM | 80 | 0.1891 ± 0.0358 | 0.7517 |

Table 3: Performance of training different cancer types jointly incorporating multi-omics data plus adding the metadata of morphological classification and anatomical locations.

| | | Gene expression+DNA methylation+CNV+VCF | |
|---|---|---|---|
| | Number of samples | RMSE | C-index |
| TCGA-ACC | 80 | 0.1389 ± 0.0193 | 0.6838 |
| TCGA-BLCA | 407 | 0.1123 ± 0.0126 | 0.7489 |
| TCGA-BRCA | 1044 | 0.0101 ± 0.0001 | 0.8620 |
| TCGA-CESC | 294 | 0.1083 ± 0.0117 | 0.8076 |
| TCGA-CHOL | 36 | 0.4271 ± 0.1824 | 0.6592 |
| TCGA-COAD | 433 | 0.0949 ± 0.0090 | 0.8364 |
| TCGA-DLBC | 37 | 0.2838 ± 0.0805 | 0.6849 |
| TCGA-ESCA | 184 | 0.1716 ± 0.0294 | 0.7747 |
| TCGA-GBM | 166 | 0.0147 ± 0.0002 | 0.8055 |
| TCGA-HNSC | 510 | 0.0282 ± 0.0008 | 0.8578 |
| TCGA-KICH | 66 | 0.1887 ± 0.0356 | 0.7351 |
| TCGA-KIRC | 339 | 0.0376 ± 0.0014 | 0.8547 |
| TCGA-KIRP | 288 | 0.0884 ± 0.0078 | 0.7789 |
| TCGA-LAML | 140 | 0.3927 ± 0.1542 | 0.6778 |
| TCGA-LGG | 511 | 0.2277 ± 0.0518 | 0.7293 |
| TCGA-LIHC | 371 | 0.2141 ± 0.0458 | 0.7153 |
| TCGA-LUAD | 509 | 0.0106 ± 0.0001 | 0.8728 |
| TCGA-LUSC | 496 | 0.0350 ± 0.0012 | 0.8643 |
| TCGA-MESO | 83 | 0.1964 ± 0.0386 | 0.7212 |
| TCGA-OV | 443 | 0.0446 ± 0.0020 | 0.8257 |
| TCGA-PAAD | 178 | 0.2172 ± 0.0472 | 0.7137 |
| TCGA-PCPG | 179 | 0.0613 ± 0.0038 | 0.8332 |
| TCGA-PRAD | 498 | 0.2875 ± 0.0827 | 0.6948 |
| TCGA-READ | 158 | 0.2004 ± 0.0402 | 0.7244 |
| TCGA-SARC | 255 | 0.0252 ± 0.0006 | 0.8539 |
| TCGA-SKCM | 456 | 0.2848 ± 0.0811 | 0.6904 |
| TCGA-STAD | 414 | 0.0684 ± 0.0047 | 0.8126 |
| TCGA-TGCT | 134 | 0.2048 ± 0.0419 | 0.6443 |
| TCGA-THCA | 496 | 0.0262 ± 0.0007 | 0.8588 |
| TCGA-THYM | 123 | 0.1875 ± 0.0352 | 0.6748 |
| TCGA-UCEC | 542 | 0.1149 ± 0.0132 | 0.7513 |
| TCGA-UCS | 55 | 0.3894 ± 0.1516 | 0.6626 |
| TCGA-UVM | 80 | 0.2574 ± 0.0663 | 0.7101 |

Table 4: Performance comparison on training different cancer types jointly incorporating multi-omics data only.

| Cancer types | Prognostic biomarker genes | Average rank |
|---|---|---|
| Breast Invasive Carcinoma (TCGA-BRCA) | PIK3CA | 7 |
| Adrenocortical Carcinoma (TCGA-ACC) | CTNNB1 | 14 |
| | TP53 | 9 |
| | ZNRF3 | 38 |
| Bladder Urothelial Carcinoma (TCGA-BLCA) | ATM | 22 |
| | ERBB3 | 16 |
| | ERCC2 | 54 |
| | FANCC | 32 |
| | RB1 | 68 |
| Cervical Squamous Cell Carcinoma and Endocervical Adenocarcinoma (TCGA-CESC) | HRAS | 20 |
| | KRAS | 6 |
| Cholangiocarcinoma (TCGA-CHOL) | IDH1 | 74 |
| | IDH2 | 63 |
| Colon Adenocarcinoma (TCGA-COAD) | APC | 184 |
| | ATM | 42 |
| | BRAF | 15 |
| | EGFR | 117 |
| | RNF43 | 78 |
| | ZNRF3 | 26 |
| | ERBB2 | 6 |
| | FBXW7 | 23 |
| | MET | 273 |
| Lymphoid Neoplasm Diffuse Large B-cell Lymphoma (TCGA-DLBC) | CD79B | 56 |
| | EZH2 | 44 |
| Esophageal Carcinoma (TCGA-ESCA) | PTEN | 10 |
| | EGFR | 136 |
| | ERBB2 | 47 |
| Glioblastoma Multiforme (TCGA-GBM) | EGFR | 289 |
| | TERT | 51 |
| Head and Neck Squamous Cell Carcinoma (TCGA-HNSC) | EGFR | 163 |
| | PIK3CB | 8 |
| | TP53 | 13 |
| | ERBB2 | 68 |
| | FGFR1 | 26 |
| Kidney Chromophobe (TCGA-KICH) | BAP1 | 5 |
| | VHL | 19 |
| Kidney Renal Clear Cell Carcinoma (TCGA-KIRC) | BAP1 | 68 |
| | VHL | 29 |
| Kidney Renal Papillary Cell Carcinoma (TCGA-KIRP) | BAP1 | 47 |
| | VHL | 13 |
| Acute Myeloid Leukemia (TCGA-LAML) | DNMT3A | 159 |
| | HRAS | 94 |
| | KIT | 56 |
| | KRAS | 20 |
| | NOTCH2 | 146 |
| | NPM1 | 34 |
| | NRAS | 83 |
| Brain Lower Grade Glioma (TCGA-LGG) | ALK | 44 |
| | ATM | 89 |
| | ATR | 257 |
| | BRAF | 63 |
| | CDKN2A | 430 |
| | CDKN2B | 424 |
| | CDKN2C | 275 |
| | EGFR | 28 |
| | IDH1 | 96 |
| | NF1 | 54 |
| | PIK3CA | 9 |
| | PIK3R1 | 46 |
| | PTEN | 5 |
| | STAG2 | 18 |
| Liver Hepatocellular Carcinoma (TCGA-LIHC) | AFP | 65 |
| Lung Adenocarcinoma (TCGA-LUAD) | ARAF | 13 |
| | BRAF | 64 |
| | ERBB2 | 95 |
| | NRAS | 16 |
| | STK11 | 168 |
| Lung Squamous Cell Carcinoma (TCGA-LUSC) | DDR2 | 52 |
| | EPHA2 | 16 |
| | FGFR2 | 276 |
| | FGFR1 | 183 |
| Mesothelioma (TCGA-MESO) | BAP1 | 24 |
| Ovarian Serous Cystadenocarcinoma (TCGA-OV) | AKT1 | 48 |
| | ATR | 365 |
| | BRAF | 94 |
| | BRCA1 | 54 |
| | BRCA2 | 60 |
| | ERCC4 | 16 |
| | ERCC6 | 22 |
| | MAP2K1 | 479 |
| | PTEN | 6 |
| | SMARCA4 | 261 |
| | TP53 | 24 |
| | ERBB2 | 11 |
| Pancreatic Adenocarcinoma (TCGA-PAAD) | BRCA1 | 146 |
| | BRCA2 | 172 |
| Pheochromocytoma and Paraganglioma (TCGA-PCPG) | HRAS | 64 |
| | TP53 | 26 |
| | KMT2D | 382 |
| | SDHA | 165 |
| | SDHC | 77 |
| | SDHAF2 | 41 |
| | FH | 37 |
| | KIF1B | 162 |
| | TMEM127 | 34 |
| Prostate Adenocarcinoma (TCGA-PRAD) | ATM | 76 |
| | BRCA1 | 25 |
| | BRCA2 | 95 |
| | CHEK2 | 270 |
| | FANCA | 126 |
| | HDAC2 | 57 |
| | PALB2 | 62 |
| | AR | 136 |
| | AURKA | 98 |
| | MYC | 345 |
| Rectum Adenocarcinoma (TCGA-READ) | KRAS | 12 |
| | TP53 | 9 |
| Sarcoma (TCGA-SARC) | MDM4 | 26 |
| Skin Cutaneous Melanoma (TCGA-SKCM) | BAP1 | 42 |
| | BRAF | 14 |
| | CDKN2A | 296 |
| | ERBB4 | 155 |
| | KIT | 37 |
| | NF1 | 63 |
| | PDGFRA | 10 |
| | CCND1 | 28 |
| | PAK1 | 75 |
| Stomach Adenocarcinoma (TCGA-STAD) | ATM | 6 |
| | FGFR2 | 24 |
| | MET | 66 |
| Testicular Germ Cell Tumors (TCGA-TGCT) | KIT | 38 |
| | KRAS | 6 |
| | NRAS | 14 |
| Thyroid Carcinoma (TCGA-THCA) | BRAF | 89 |
| | PIK3CA | 11 |
| | PTEN | 6 |
| | RET | 274 |
| | TSC2 | 54 |
| Thymoma (TCGA-THYM) | KIT | 195 |
| | TP53 | 26 |
| Uterine Corpus Endometrial Carcinoma (TCGA-UCEC) | PTEN | 7 |
| | PIK3CA | 15 |
| | TP53 | 13 |
| Uterine Carcinosarcoma (TCGA-UCS) | TP53 | 27 |
| | FBXW7 | 125 |
| | PIK3CA | 39 |
| Uveal Melanoma (TCGA-UVM) | BAP1 | 43 |
| | EIF1AX | 376 |
| | SF3B1 | 91 |



Table 5: Average rank results for prognostic markers based on the attention mechanism

| Cancer Type | Description | Overall samples count | Samples with survival data | Gene expression | DNA methylation | CNVs | SNVs | Clinical data | Used samples count |
|---|---|---|---|---|---|---|---|---|---|
| TCGA-ACC | Adrenocortical Carcinoma | 92 | 91 | 80 | 80 | 92 | 92 | 92 | 80 |
| TCGA-BLCA | Bladder Urothelial Carcinoma | 412 | 407 | 412 | 412 | 412 | 412 | 412 | 407 |
| TCGA-BRCA | Breast Invasive Carcinoma | 1098 | 1076 | 1097 | 1095 | 1098 | 1044 | 1098 | 1044 |
| TCGA-CESC | Cervical Squamous Cell Carcinoma and Endocervical Adenocarcinoma | 307 | 294 | 307 | 307 | 304 | 305 | 307 | 294 |
| TCGA-CHOL | Cholangiocarcinoma | 51 | 48 | 36 | 36 | 36 | 51 | 51 | 36 |
| TCGA-COAD | Colon Adenocarcinoma | 461 | 437 | 459 | 458 | 460 | 433 | 461 | 433 |
| TCGA-DLBC | Lymphoid Neoplasm Diffuse Large B-cell Lymphoma | 58 | 47 | 48 | 48 | 50 | 37 | 58 | 37 |
| TCGA-ESCA | Esophageal Carcinoma | 185 | 184 | 184 | 185 | 184 | 185 | 185 | 184 |
| TCGA-GBM | Glioblastoma Multiforme | 617 | 595 | 166 | 423 | 599 | 396 | 617 | 166 |
| TCGA-HNSC | Head and Neck Squamous Cell Carcinoma | 528 | 527 | 528 | 528 | 526 | 510 | 528 | 510 |
| TCGA-KICH | Kidney Chromophobe | 113 | 112 | 66 | 66 | 66 | 66 | 113 | 66 |
| TCGA-KIRC | Kidney Renal Clear Cell Carcinoma | 537 | 533 | 534 | 533 | 534 | 339 | 537 | 339 |
| TCGA-KIRP | Kidney Renal Papillary Cell Carcinoma | 291 | 288 | 291 | 291 | 290 | 288 | 291 | 288 |
| TCGA-LAML | Acute Myeloid Leukemia | 200 | 175 | 188 | 140 | 200 | 149 | 200 | 140 |
| TCGA-LGG | Brain Lower Grade Glioma | 516 | 511 | 516 | 516 | 515 | 513 | 516 | 511 |
| TCGA-LIHC | Liver Hepatocellular Carcinoma | 377 | 371 | 376 | 377 | 376 | 375 | 377 | 371 |
| TCGA-LUAD | Lung Adenocarcinoma | 585 | 509 | 519 | 579 | 518 | 569 | 585 | 509 |
| TCGA-LUSC | Lung Squamous Cell Carcinoma | 504 | 496 | 504 | 503 | 504 | 497 | 504 | 496 |
| TCGA-MESO | Mesothelioma | 87 | 85 | 87 | 87 | 83 | 87 | 87 | 83 |
| TCGA-OV | Ovarian Serous Cystadenocarcinoma | 608 | 583 | 492 | 602 | 497 | 443 | 608 | 443 |
| TCGA-PAAD | Pancreatic Adenocarcinoma | 185 | 184 | 178 | 184 | 185 | 183 | 185 | 178 |
| TCGA-PCPG | Pheochromocytoma and Paraganglioma | 179 | 179 | 179 | 179 | 179 | 179 | 179 | 179 |
| TCGA-PRAD | Prostate Adenocarcinoma | 500 | 500 | 498 | 498 | 498 | 498 | 498 | 498 |
| TCGA-READ | Rectum Adenocarcinoma | 172 | 162 | 167 | 165 | 167 | 158 | 172 | 158 |
| TCGA-SARC | Sarcoma | 261 | 258 | 261 | 261 | 261 | 255 | 261 | 255 |
| TCGA-SKCM | Skin Cutaneous Melanoma | 470 | 456 | 469 | 470 | 470 | 470 | 470 | 456 |
| TCGA-STAD | Stomach Adenocarcinoma | 443 | 414 | 439 | 443 | 443 | 441 | 443 | 414 |
| TCGA-TGCT | Testicular Germ Cell Tumors | 150 | 134 | 150 | 150 | 150 | 150 | 150 | 134 |
| TCGA-THCA | Thyroid Carcinoma | 507 | 506 | 507 | 507 | 505 | 496 | 507 | 496 |
| TCGA-THYM | Thymoma | 124 | 123 | 124 | 124 | 124 | 123 | 124 | 123 |
| TCGA-UCEC | Uterine Corpus Endometrial Carcinoma | 560 | 544 | 559 | 559 | 558 | 542 | 560 | 542 |
| TCGA-UCS | Uterine Carcinosarcoma | 57 | 55 | 57 | 57 | 57 | 57 | 57 | 55 |
| TCGA-UVM | Uveal Melanoma | 80 | 80 | 80 | 80 | 80 | 80 | 80 | 80 |
| The total | | 11,315 | 10,964 | 10,558 | 10,943 | 11,126 | 10,418 | 11,315 | 10,005 |

Table 6: Summary of TCGA data used in our work.

| Cancer types | Morphological classification |
|---|---|
| Breast Invasive Carcinoma (TCGA-BRCA) | Epithelial Neoplasm |
| Adrenocortical Carcinoma (TCGA-ACC) | Epithelial Neoplasm |
| Bladder Urothelial Carcinoma (TCGA-BLCA) | Epithelial Neoplasm |
| Cervical Squamous Cell Carcinoma and Endocervical Adenocarcinoma (TCGA-CESC) | Epithelial Neoplasm |
| Cholangiocarcinoma (TCGA-CHOL) | Epithelial Neoplasm |
| Colon Adenocarcinoma (TCGA-COAD) | Epithelial Neoplasm |
| Lymphoid Neoplasm Diffuse Large B-cell Lymphoma (TCGA-DLBC) | Hematopoietic and Lymphoid Cell Neoplasm |
| Esophageal Carcinoma (TCGA-ESCA) | Epithelial Neoplasm |
| Glioblastoma Multiforme (TCGA-GBM) | Neuroepithelial, Perineurial, and Schwann Cell Neoplasm |
| Head and Neck Squamous Cell Carcinoma (TCGA-HNSC) | Epithelial Neoplasm |
| Kidney Chromophobe (TCGA-KICH) | Epithelial Neoplasm |
| Kidney Renal Clear Cell Carcinoma (TCGA-KIRC) | Epithelial Neoplasm |
| Kidney Renal Papillary Cell Carcinoma (TCGA-KIRP) | Epithelial Neoplasm |
| Acute Myeloid Leukemia (TCGA-LAML) | Hematopoietic and Lymphoid Cell Neoplasm |
| Brain Lower Grade Glioma (TCGA-LGG) | Neuroepithelial, Perineurial, and Schwann Cell Neoplasm |
| Liver Hepatocellular Carcinoma (TCGA-LIHC) | Epithelial Neoplasm |
| Lung Adenocarcinoma (TCGA-LUAD) | Epithelial Neoplasm |
| Lung Squamous Cell Carcinoma (TCGA-LUSC) | Epithelial Neoplasm |
| Mesothelioma (TCGA-MESO) | Mesothelial Neoplasm |
| Ovarian Serous Cystadenocarcinoma (TCGA-OV) | Epithelial Neoplasm |
| Pancreatic Adenocarcinoma (TCGA-PAAD) | Epithelial Neoplasm |
| Pheochromocytoma and Paraganglioma (TCGA-PCPG) | Neural Crest Cell Tumors |
| Prostate Adenocarcinoma (TCGA-PRAD) | Epithelial Neoplasm |
| Rectum Adenocarcinoma (TCGA-READ) | Epithelial Neoplasm |
| Sarcoma (TCGA-SARC) | Mesenchymal Cell Neoplasm |
| Skin Cutaneous Melanoma (TCGA-SKCM) | Neural Crest Cell Tumors |
| Stomach Adenocarcinoma (TCGA-STAD) | Epithelial Neoplasm |
| Testicular Germ Cell Tumors (TCGA-TGCT) | Germ Cell Tumor |
| Thyroid Carcinoma (TCGA-THCA) | Epithelial Neoplasm |
| Thymoma (TCGA-THYM) | Epithelial Neoplasm |
| Uterine Corpus Endometrial Carcinoma (TCGA-UCEC) | Epithelial Neoplasm |
| Uterine Carcinosarcoma (TCGA-UCS) | Epithelial Neoplasm |
| Uveal Melanoma (TCGA-UVM) | Neural Crest Cell Tumors |

Table 7: Mapping of cancer types to their morphological classification.

| Cancer types | Anatomical locations |
|---|---|
| Breast Invasive Carcinoma (TCGA-BRCA) | Breast |
| Adrenocortical Carcinoma (TCGA-ACC) | Adrenal gland |
| Bladder Urothelial Carcinoma (TCGA-BLCA) | Bladder |
| Cervical Squamous Cell Carcinoma and Endocervical Adenocarcinoma (TCGA-CESC) | Cervix uteri |
| Cholangiocarcinoma (TCGA-CHOL) | Gallbladder |
| | Liver and intrahepatic bile ducts |
| Colon Adenocarcinoma (TCGA-COAD) | Colon |
| | Rectosigmoid junction |
| Lymphoid Neoplasm Diffuse Large B-cell Lymphoma (TCGA-DLBC) | Bones, joints and articular cartilage of other and unspecified sites |
| | Brain |
| | Breast |
| | Colon |
| | Connective, subcutaneous and other soft tissues |
| | Heart, mediastinum, and pleura |
| | Hematopoietic and reticuloendothelial systems |
| | Lymph nodes |
| | Other and unspecified major salivary glands |
| | Retroperitoneum and peritoneum |
| | Small intestine |
| | Stomach |
| | Testis |
| | Thyroid gland |
| Esophageal Carcinoma (TCGA-ESCA) | Esophagus |
| | Stomach |
| Glioblastoma Multiforme (TCGA-GBM) | Brain |
| Head and Neck Squamous Cell Carcinoma (TCGA-HNSC) | Base of tongue |
| | Bones, joints and articular cartilage of other and unspecified sites |
| | Floor of mouth |
| | Gum |
| | Hypopharynx |
| | Larynx |
| | Lip |
| | Oropharynx |
| | Other and ill-defined sites in lip, oral cavity and pharynx |
| | Other and unspecified parts of mouth |
| | Other and unspecified parts of tongue |
| | Palate |
| | Tonsil |
| Kidney Chromophobe (TCGA-KICH) | Kidney |
| Kidney Renal Clear Cell Carcinoma (TCGA-KIRC) | Kidney |
| Kidney Renal Papillary Cell Carcinoma (TCGA-KIRP) | Kidney |
| Acute Myeloid Leukemia (TCGA-LAML) | Hematopoietic and reticuloendothelial systems |
| Brain Lower Grade Glioma (TCGA-LGG) | Brain |
| Liver Hepatocellular Carcinoma (TCGA-LIHC) | Liver and intrahepatic bile ducts |
| Lung Adenocarcinoma (TCGA-LUAD) | Bronchus and lung |
| Lung Squamous Cell Carcinoma (TCGA-LUSC) | Bronchus and lung |
| Mesothelioma (TCGA-MESO) | Bronchus and lung |
| | Heart, mediastinum, and pleura |
| Ovarian Serous Cystadenocarcinoma (TCGA-OV) | Ovary |
| Pancreatic Adenocarcinoma (TCGA-PAAD) | Pancreas |
| Pheochromocytoma and Paraganglioma (TCGA-PCPG) | Adrenal gland |
| | Heart, mediastinum, and pleura |
| | Other and ill-defined sites |
| | Other endocrine glands and related structures |
| | Retroperitoneum and peritoneum |
| | Spinal cord, cranial nerves, and other parts of central nervous system |
| Prostate Adenocarcinoma (TCGA-PRAD) | Prostate gland |
| Rectum Adenocarcinoma (TCGA-READ) | Colon |
| | Rectosigmoid junction |
| | Rectum |
| Sarcoma (TCGA-SARC) | Bones, joints and articular cartilage of limbs |
| | Colon |
| | Connective, subcutaneous and other soft tissues |
| | Corpus uteri |
| | Kidney |
| | Meninges |
| | Other and unspecified male genital organs |
| | Other and unspecified parts of tongue |
| | Ovary |
| | Peripheral nerves and autonomic nervous system |
| | Retroperitoneum and peritoneum |
| | Stomach |
| | Uterus, NOS |
| Skin Cutaneous Melanoma (TCGA-SKCM) | Skin |
| Stomach Adenocarcinoma (TCGA-STAD) | Stomach |
| Testicular Germ Cell Tumors (TCGA-TGCT) | Testis |
| Thyroid Carcinoma (TCGA-THCA) | Thyroid gland |
| Thymoma (TCGA-THYM) | Thymus |
| | Heart, mediastinum, and pleura |
| Uterine Carcinosarcoma (TCGA-UCS) | Uterus, NOS |
| Uveal Melanoma (TCGA-UVM) | Eye and adnexa |

Table 8: Anatomical locations observed for each cancer type within TCGA.

| | Breast | | Lung | | Glioblastoma | |
|---|---|---|---|---|---|---|
| | RMSE | C-index | RMSE | C-index | RMSE | C-index |
| GCNConv | **0.0101** | **0.8593** | **0.0111** | **0.8663** | 0.0159 | 0.7941 |
| GENConv | 0.0253 | 0.8242 | 0.0380 | 0.7834 | **0.0108** | **0.8082** |
| GraphSAGE | 0.0311 | 0.8149 | 0.0423 | 0.7916 | 0.0488 | 0.7752 |
| GraphConv | 0.0498 | 0.7549 | 0.0258 | 0.7595 | 0.0338 | 0.7053 |
| APPNP | 0.0349 | 0.8038 | 0.0237 | 0.7684 | 0.0315 | 0.7186 |

Table 9: Performance comparison of the Graph Convolutional Network with other graph-based neural network architectures trained individually on Breast, Lung and Glioblastoma.
